# Parallel multicopy-suppressor screens reveal convergent evolution of phage-encoded single gene lysis proteins

**DOI:** 10.1101/2022.01.20.477139

**Authors:** Benjamin A. Adler, Karthik Chamakura, Heloise Carion, Jonathan Krog, Adam M. Deutschbauer, Ryland F Young, Vivek K. Mutalik, Adam P. Arkin

## Abstract

In contrast to dsDNA phages where multiple proteins are involved in programmed host lysis, lysis in ssRNA *Fiersviridae* and ssDNA *Microviridae* phages requires only a single gene (*sgl for <u>s</u>ingle <u>g</u>ene <u>l</u>ysis*) to meet the size constraints of some of the smallest genomes in the biosphere. To achieve lysis, Sgl proteins exploit evolutionary “weak spots” in bacterial cell wall biogenesis. In several cases, this is done by inhibiting specific steps in Lipid II synthesis. Recently metatranscriptomics has revealed thousands of novel ssRNA phage genomes, each of which must carry at least one *sgl* gene. Determining the targets of these Sgl proteins could reveal novel vulnerabilities in bacterial envelope biogenesis and may lead to new antibiotics. Here, we employ a high-throughput genetic screen to uncover genome-wide host suppressors of Sgl activity and apply it to a set of diverse Sgls with unknown molecular targets. In addition to validating known molecular mechanisms, we determined that the Sgl of PP7, an ssRNA phage of *P. aeruginosa*, targets MurJ, the flippase responsible for Lipid II export which was previously shown to be the target of the Sgl of coliphage M. These two Sgls, which are unrelated and predicted to have opposite membrane topology, thus represent a case of convergent evolution. Another set of Sgls which are thought to cause lysis without inhibiting cell wall synthesis elicit a common set of multicopy suppressors, suggesting these Sgls act by the same or similar mechanism.

## Introduction

Lysis of the bacterial host is the last step in the bacteriophage (phage) life cycle, determined by the lytic program encoded on the phage genome. In double-strand DNA (dsDNA) phages, lysis is mediated by multiprotein systems that disrupt the cytoplasmic/inner membrane, degrade the peptidoglycan/cell wall, compromise the outer membrane, and regulate the lytic process^1^. In contrast, single strand RNA (ssRNA) phages and lytic single-strand DNA (ssDNA) phages use the product of a single gene (*sgl*) to carry out host lysis ^2,3^.

Until recently, only 11 *sgl*s had been identified. Of these 11, the Sgls of the coliphages ΦX174, Qβ and M were shown to block the production and translocation of periplasmic Lipid II, the universal precursor for peptidoglycan (PG) synthesis, by inhibiting the conserved enzymes MraY, MurA and MurJ, respectively (Fig 1a). The situation is mysterious, however, for the canonical male-specific coliphage MS2, which in 1975 was the first genetic entity to have its complete genome published^4^. Mutational analysis revealed that a cryptic 75 codon reading frame was required for lysis^5^. This Sgl, named L, caused lysis when expressed alone from a plasmid vector; subsequent studies reported it to be a membrane protein^6^ and to support lysis without inhibiting net peptidoglycan (PG) biosynthesis, as measured by incorporation of ^3^H-mDAP^7^. More recent genetic analysis has shown that L function requires formation of a complex with the chaperone DnaJ^8^. Moreover, mutational analysis and comparison with other Sgls led to the hypothesis that the other 7 known Sgls were “L-like”, in that despite the lack of sequence similarity, they shared a characteristic 4 domain structure, including a characteristic Leu-Ser motif at the C-terminus of a hydrophobic domain^9^. On this basis, it was proposed that the L-like Sgl family shared a common target, conserved in their diverse bacterial hosts (Pseudomonas, Acinetobacter, Caulobacter, and *E. coli*).

**Fig. 1:**
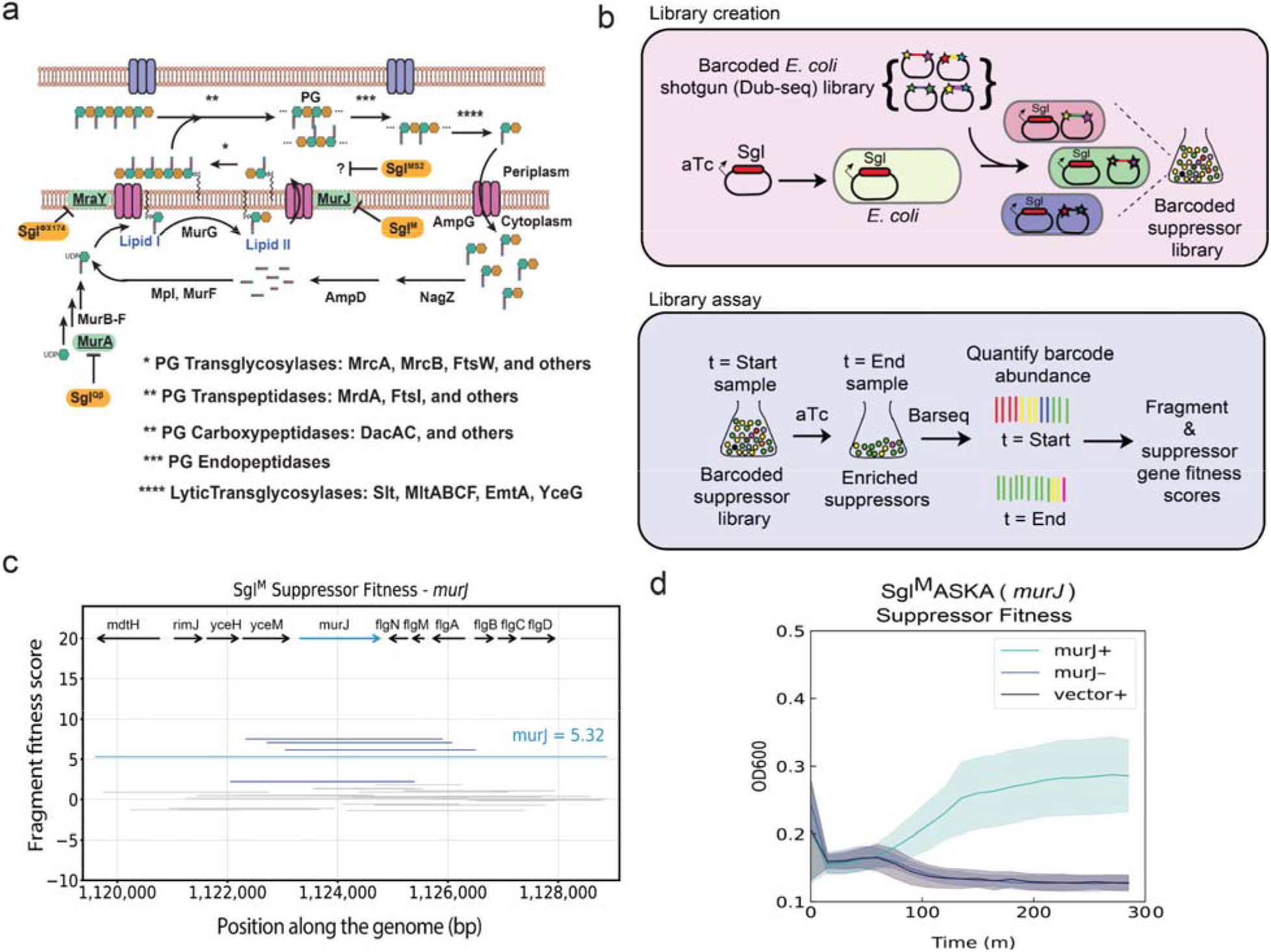
Genome-wide screen to identify host suppressors of phage-encoded single gene lysis systems. **(a)** Schematic representation of PG maturation and recycling in *E. coli* **^20^** with a focus on genetic interactions observed in the Sgl protein literature and this study. Three well-studied Sgl proteins (orange) are shown with their primary targets (green). In brief, the PG precursor Lipid II is synthesized through the serial activity of enzymes MurA-F, MraY, and MurG, and then translocated to the periplasm via the flippase MurJ. The disaccharide oligopeptide moiety of Lipid II is then added to the PG, crosslinked, and tailored with other PG polymers through coordinated activity of PG-transglycosylases (*), PG-transpeptidases (**), and PG-carboxypeptidases (**) to form a mature PG. Mature PG is broken down into recyclable monomers through endopeptidase (***) and lytic transglycosylase (****) activities. Such monomers can be recycled through AmpG import, further breakdown by NagZ and AmpD (and other enzymes not shown), and re-ligation to the substrate for MurF by Mpl. Additional regulation on PG maturation is imposed by the rod and division networks and discussed in detail elsewhere ^21^. (b) Cartoon description of suppressor library creation and assay. A toxin gene (in this paper, an *sgl* gene) is cloned into an aTc-inducible vector. The Dub-seq library ^15^ containing previously mapped dual barcoded (shown as stars on the plasmid) random genomic fragments (shown as colored regions between stars) from BW25113 *E. coli* strain is transformed into the DH10B *E. coli* strain carrying the cloned *sgl* gene ^15^, creating a barcoded suppressor library consisting of tens of thousands of independent transformants. This library was grown to OD600 ~1.0, where a “t = start” sample representing an initial unbiased distribution of genomic fragments, was collected. We then induced the cultures with aTc, biasing the library towards suppressing fragments. Strains were tracked by quantifying the abundance of DNA barcodes associated with each strain by Illumina sequencing. Sgl-specific gene fitness profiles were calculated by taking the log2-fold-change of barcode abundances between post- (t=End) and pre- (t=Start) induction of *sgl* and fragment and gene fitness scores calculated as described in methods. (c) Representative fragment and gene fitness data from our suppressor screening experiment for Sgl^M^. Dub-seq fragment (strain) data (*y* axis) for the genomic region (*x* axis) surrounding *murJ* under induction of Sgl^M^ is shown. Each purple and gray line is a Dub-seq fragment. Those that completely cover *murJ* are shown in purple and fragments that do not contain *murJ* or cover partially are colored gray. The *murJ* gene fitness score of 5.32, estimated using a regression model, is shown as a blue line (Methods)^15^.Multiple barcodes representing fragments containing MurJ were specifically enriched in our Sgl^M^ screens. (d) Growth curves show heterologous expression of wildtype MurJ can suppress Sgl^M^ lytic activity. Teal represents *sgl^M^* and *murJ* co-overexpression using 16.1 ng/μL aTc and 25μM IPTG respectively. Blue represents *sgl^M^* expression in the absence of *murJ* induction using 16.1 ng/μL aTc without IPTG. Black represents *sgl^M^* expression in the presence of an empty ASKA vector using 16.1 ng/μL aTc and 25μM IPTG. All growth curves represent 3 biological replicates.

Interest in the Sgl field suddenly increased when, beginning in 2016, environmental metagenome and transcriptome mining increased the available sequence diversity of *Fiersviridae* to upwards of 10,000 genomes^10–12^. Although the hosts of these ssRNA phages are unknown, each genome is expected to encode at least one Sgl. The prospect of identifying thousands of diverse Sgls, each with the capability of finding a “weak spot” in bacterial biogenesis, is alluring, not the least because it might lead to opportunities for antibiotic development. However, the classic molecular genetic approach has required decades to find three Sgl targets and has left the L mechanism still enigmatic. Thus, unbiased and scalable screening platforms may be required to realize the promise and diversity of Sgls ^13,14^. One approach to this problem may be Dub-seq, which is founded on doubly barcoded over-expression libraries of randomly sheared bacterial DNA^15^. Once the barcodes are mapped to chromosomal segments, the Dub-seq library can be used for competitive fitness assays^15^. We have demonstrated the utility, scalability and barcode standardization of fitness assays by studying the tolerance phenotypes against diverse antibiotics, stressors and metals, and most recently used to characterize genetic barriers in phage-host interactions in *E. coli* ^15,16^.

In this study, we repurpose Dub-seq for genome-wide assessment of host suppressors of Sgl activity and apply it to five diverse ssRNA Sgls awaiting molecular target characterization. We establish the screening platform by recapitulating the known molecular target of Sgl^M^. The results enable a rapid determination of one of the Sgl targets and suggest common or similar mechanisms for most of the others.

## Results

### Devising rapid Sgl suppressor identification screens

Previously Sgl targets have been identified by inducing a multicopy plasmid clone of the *sgl* and selecting for spontaneous missense mutations in the target genes^17–19^ or for multicopy suppressors using an *E. coli* gene library (Sgl^M19^). Both approaches are constrained, the former for availability of mutable sites that block Sgl/target interaction without destroying function and the latter for appropriate Sgl/target copy number and affinities. We hypothesized that expressing an *sgl* gene in the context of a barcoded shotgun expression library of the host (for example, *E. coli* Dub-seq library expressed in *E. coli*) and using barcode sequencing as a readout would enable the quick identification of all genes in a genome that contribute to fitness, including the target gene, even if Sgl/target levels were not ideal (Figs. 1b). As a proof-of-principle, we adapted our Dub-seq platform for screening suppressors against the toxicity of Sgl^M^, an inhibitor of the lipid II flippase, MurJ (Fig. 1a-d) ^19^. Sgl^M^ was chosen because the first evidence for its targeting MurJ came from a multicopy-suppression of induced *sgl^M^* lysis using an *E. coli* gene library^19^. Briefly, we cloned *sgl^M^* into a low-copy plasmid under an anhydrotetracycline (aTc)-inducible promoter and showed that induction caused lysis in *E. coli* K-12 (Fig. 1d, Methods). We then moved a previously characterized *E. coli* Dub-seq plasmid library (pFAB5516), consisting of *E. coli* genomic DNA fragments cloned between two 20bp random DNA barcodes into our assay strain DH10B *E. coli* cells ^15^. This process resulted in a library of 17,007 unique members, (BA1320L), with each strain harboring an inducible *sgl^M^* vector and a unique member of the pFAB5516 library (Fig. 1b) (Supplementary Table 1). We then subjected this BA1320L library to Sgl^M^ induction in liquid culture, isolated plasmid DNA from cells collected before and after induction and subjected the DNA to BarSeq PCR. The product was sequenced on a Hiseq4000 platform (Methods) and analyzed for the change in barcode abundance (a proxy for cloned genomic region consisting ~1-3 genes) after Sgl^M^ induction. We then calculated the fragment fitness score for each strain by taking the normalized log2 ratio of the number of reads for each barcode at the end and at the start of the experiment (Fig. 1b). Positive scores indicate that the gene(s) contained on that fragment lead to an increase in relative fitness, whereas negative values mean the gene(s) on the fragment cause reduced relative fitness. Considering each fragment may contain causative and non-causative genes, we follow a regression model that examines the score of all fragments containing the gene and computes gene fitness scores, as reported earlier (Methods) ^15^. We classified genes with a fitness score >3 as high confidence hits if they have sufficient read coverage (>25 reads/barcode for both t=0 and the experiment) and these fitness effects were consistent across multiple fragments that cover the genes and across replicate experiments (Methods). We observed that induction of Sgl^M^ yielded reproducible data (N=2 fitness experiments; Supplementary Fig. 1). As expected, *murJ* emerged as a consistently enriched gene covered by multiple fragments in our screen, (Fig. 1c, Supplementary Fig. 1); moreover, heterologous expression of wildtype *murJ* suppressed the lytic activity of induced *sgl^M^* (Fig. 1d). These experiments indicated that the Sgl-dependent growth defect coupled with Dub-seq suppressor screen could correctly identify host factors that, when overexpressed, overcome the toxicity of the Sgl protein and thus could map the Sgl target to host pathways.

### Extending Dub-seq Suppressor Screens to Additional sgl Genes

The success of using Dub-seq to identify MurJ as the target of Sgl^M^ encouraged us to apply the method to Sgl^MS2^, the L protein, which has remained mechanistically uncharacterized for nearly half a century. Moreover, we included in the test set the Sgls from four other ssRNA phages: KU1, Hgal1, PRR1 and PP7 (Fig 2a). This set represents a spectrum of diversity in terms of the cellular environment within which the Sgl must function. KU1 is F-specific and thus restricted to *E. coli* and closely related *Enterobacteriaceae*. Both Hgal1 and PRR1 use the conjugative pili of multi-drug resistance plasmids as receptors and thus must function in rather diverse host environments. In contrast, PP7 recognizes the polar pilus of *Pseudomonas*. In any case, as noted above all five Sgls in this test set are proposed to be “L-like” Sgls and thus should have the same cellular target, based on sharing the four-motif organization revealed in the mutational analysis of Sgl^MS2^ ^9^. For these five Sgls (and a control), we performed 12 genome-wide suppressor screens, collected suppressor candidates, and processed BarSeq PCR samples for deep sequencing (Methods, experimental and library overviews are described in Extended Data 1 and Extended Data 2 respectively). After curation of the dataset for sufficient read coverage and consistency, we identified 190 high-confidence hits encompassing 96 genes across the 5 suppressor Dub-seq screens (Fig 2b, Methods, complete library read counts, fragment scores, gene scores are presented in Extended Data 3-5 respectively). Thus about 2% of genes exhibited at least one suppression phenotype. There were two genes, *waaQ* and *galE*, that appeared as multicopy suppressors of all five Sgls. Overproduction of GalE, which codes for UDP-glucose-4-epimerase has been shown to provide fitness benefits in earlier genetic screens, probably playing a role in modulating outer membrane biogenesis ^15,22^. Furthermore, the assay strain we used in this work is a *galE* mutant and overexpression of WT *galE* probably provides fitness benefits to the cell under Sgl-induced toxicity. The fragments carrying *waaQ* would also produce the RirA sRNA which activates the transcription of *rpoE* encoding the sigma factor for genes involved in periplasmic and OM maintenance. Neither *waaQ* or *rpoE* knockouts have overt lytic phenotypes but have been shown to exhibit some sensitivity to detergents and a plant-based antibacterial agent^23^. Thus neither *galE* or *waaQ* is likely to be the target of any of these Sgls and, instead, are genes which when over-expressed indirectly mitigate Sgl toxicity. More striking is the number of suppressor genes identified (96 total) for a group of four Sgls: MS2 (38 genes), KU1 (36 genes), Hgal1 (55 genes) and PRR1 Sgls (55 genes). Forty-five genes suppress at least two of these four Sgls, of which 22 suppress at least three and 10 suppress all four (Extended Data 5, Supplementary Figs. 2-6). In contrast, excluding *waaQ* and *galE*, Sgl^PP7^ toxicity was suppressed by only six genes, one of which was *murJ*.

**Fig. 2:**
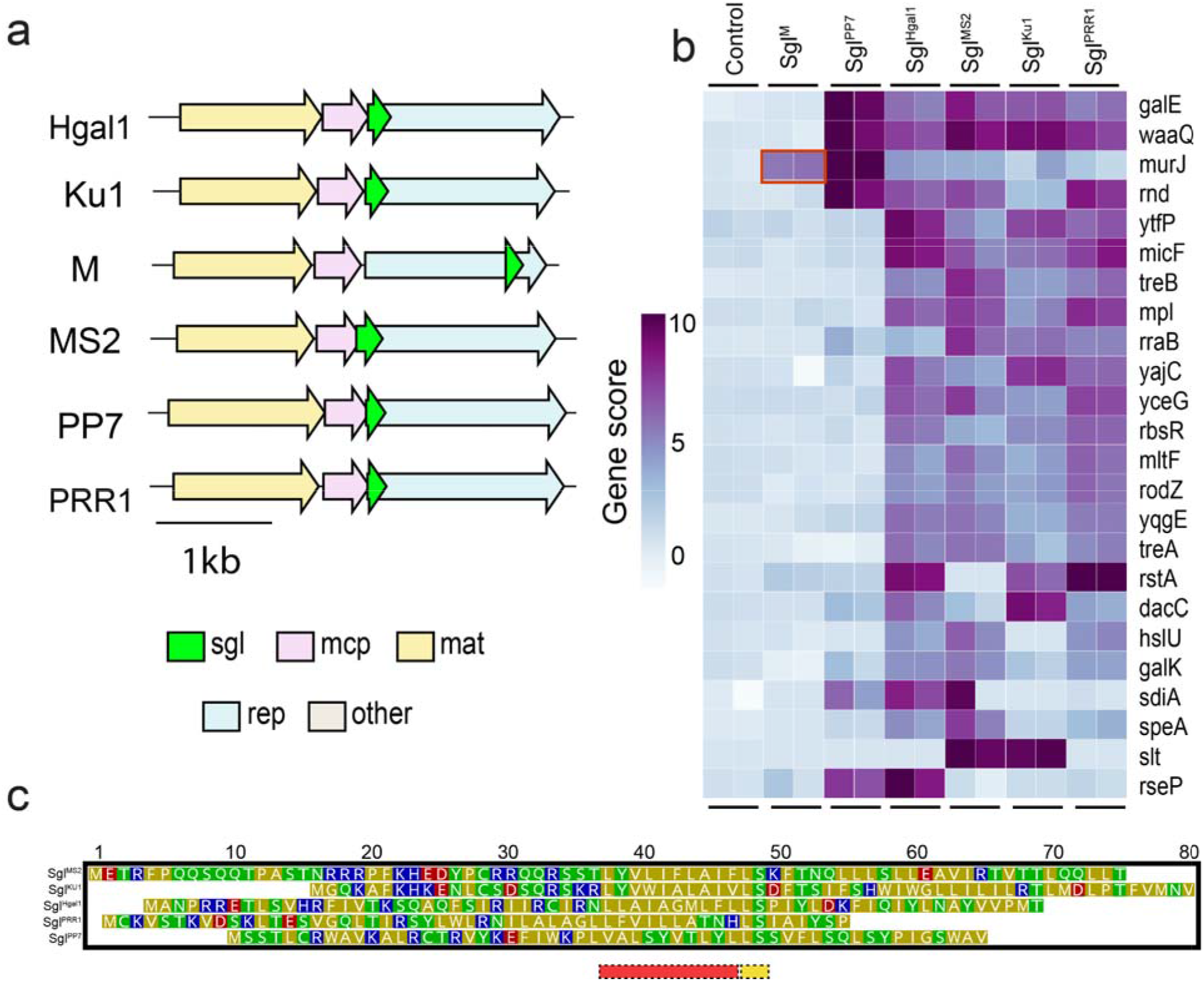
Sgl diversity in genomic context, sequence identity, and suppressor genotypes. **(a)** All *Fiersviridae-derived* Sgls investigated in this study are shown within their native genomic context. All lysis genes (green) occur in sequences overlapping with 1 or more additional genes. **(b)** Multicopy suppressors of lysis proteins as identified through high throughput gain of function screening. A curated selection of high confidence, top-scoring genes are shown for visualization purposes. Multicopy suppressors identified from prior work are boxed in red. **(c)** Sequence alignment of *Fiersviridae* lysis proteins investigated here shows that they bear little resemblance to each other. Sequence alignment was done manually(Chamakura et al. 2017). Acidic and basic residues are in red and blue respectively, while polar and nonpolar residues are shown in green and yellow respectively. The conserved LS motif **is** shown as a yellow box, preceded by a stretch of hydrophobic residues as a red box.

Although Sgl^PP7^ and Sgl^M^ shared no significant sequence similarity with each other (Fig 2c, Supplementary Fig. 7, 15.6% sequence identity, MUSCLE BLOSUM62 matrix(Edgar 2004)) but both yielded high scoring *murJ*-containing fragments (Figs. 1**e**, 3a), we wondered if MurJ could be the target of Sgl^PP7^. To validate the Mu**rJ** suppression of Sgl^PP7^, we transferred the *murJ* plasmids from the ASKA collection into *E. coli* and tested for the ability to inhibit Sgl^PP7^ lysis in liquid culture after induction with aTc (for the *sgl*) and IPTG (for the candidate target gene (Fig. 3b, Methods) ^25^. The results clearly show that the multicopy *murJ* clone from the ASKA plasmid library can block lysis in an induction-specific fashion; i.e., it is not just the presence of the multicopy gene that affects suppression. A similar result was obtained for another hit, *rseP* (Fig 3c,d), which encodes a protease known to cleave membrane proteins^26^. The simplest notion is that MurJ blocks lysis by titrating out the Sgl, whereas RseP acts to degrade it.

**Fig. 3:**
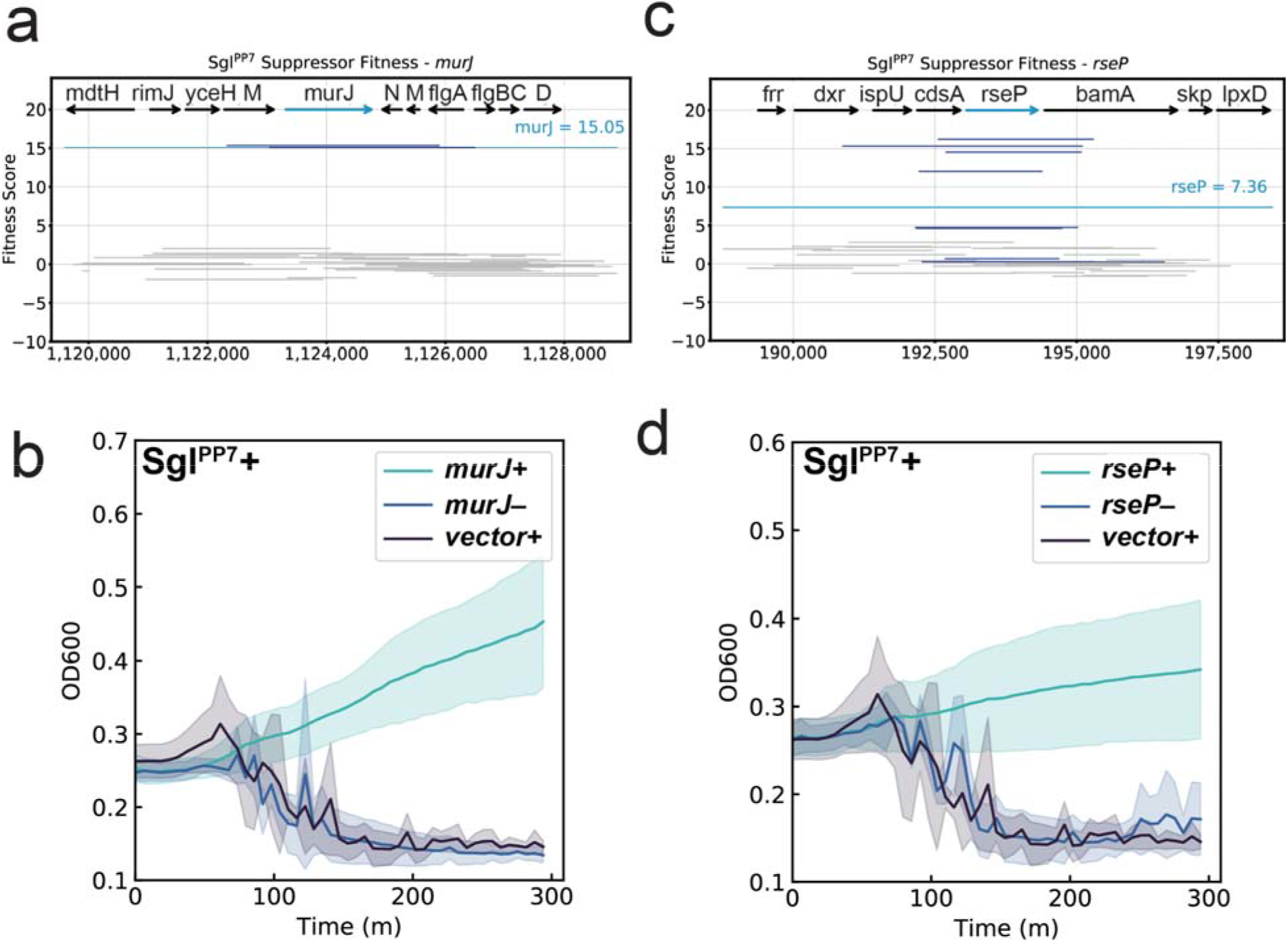
Sampling of validated multicopy suppressors against Sgl protein expression. **(a,b)** Panels refer to suppressor activity of *murJ* expression against Sgl^PP7^-mediated lysis. **(c,d)** Panels refer to suppressor activity of *rseP* expression against Sgl^PP7^-mediated lysis. Dub-seq fragment plots for the highlighted suppressor locus are shown in panels (a,c). Dark blue lines correspond to fragments covering the gene of interest. Gray lines correspond to fragments not covering the gene of interest. Teal line corresponds to gene fitness score. (b,d) Panels show lysis inhibition growth effects from a 96-well microplate reader assay. Multicopy suppressors or empty vector controls were expressed from the corresponding ASKA mutant collection plasmid under IPTG control. Teal curves correspond to suppressor induction at 50 μM IPTG. Black curves correspond to the empty ASKA vector negative control. Blue curves correspond to the uninduced suppressor plasmid. All curves are plotted as mean of three biological replicates with interpolated ± standard deviation confidence intervals. Large variations in optical density are caused by the aggregation of viscous cell debris observed in the course of the microplate reader experiment.

To further investigate the activity and specificity of MurJ suppression of Sgl^PP7^-induced lysis, we co-expressed Sgl^PP7^ with the heterologous lipid II flippases MurJ_TA_ from *Thermsipho africanus* or Amj from *Bacillus subtilis* (Fig. 4a,b). The results indicate that Sgl^PP7^ lethality can be rescued by expression of heterologous lipid II flippases,which strongly suggests that Sgl^PP7^ targets MurJ. Moreover, we obtained unambiguous evidence for the Sgl^PP7^-MurJ interaction using a genetic approach. We constructed a fusion gene, *gfp-sgl^PP7,^* that exhibited enhanced lytic function (Fig. 4b), allowing us to select spontaneous mutants that survived the induction of the fusion gene (Fig. 4b). Analysis of the survivors revealed a single amino acid substitution in MurJ conferring Sgl^PP7^-resistance: Q244P (Fig 4c). This missense change is localized to transmembrane domain 7 (TMD7), one of the 14 transmembrane domains that define the solvent-exposed cavity of MurJ and, specifically, undergo a major conformation shift as MurJ alternates between cytoplasmic- and periplasmic-open states^19,27^. Considering this amino acid change was previously observed to confer resistance to Sgl^M^, these two dissimilar proteins may target the same molecular interface of MurJ ^19^.

**Fig. 4:**
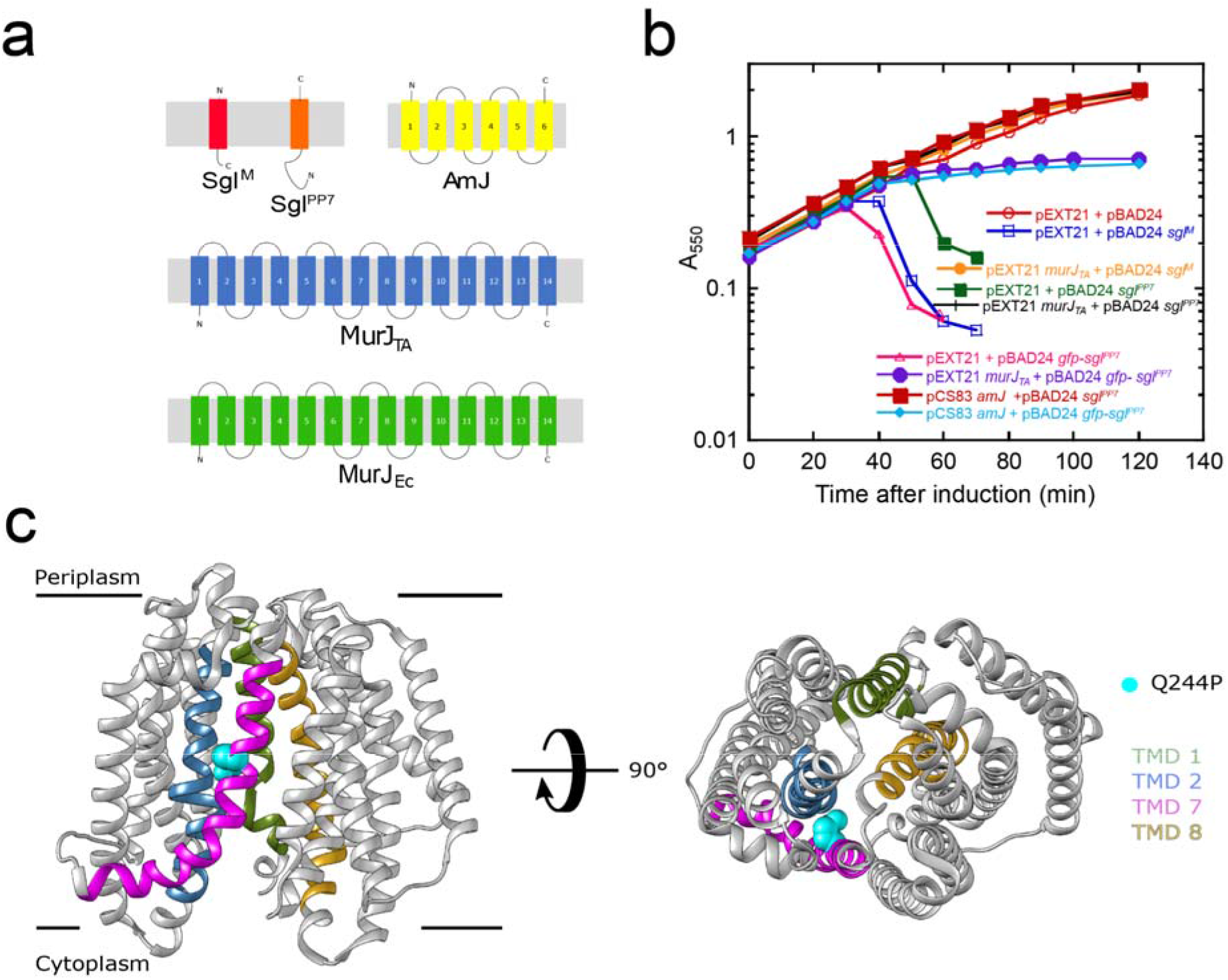
Sgl^PP7^-MurJ interaction. **(a)** Predicted membrane topologies of Sgl^M^ (red), Sgl^PP7^ (orange), AmJ (yellow), MurJ_TA_ (*Thermosipho africanus*) (blue), and MurJ_EC_ (*Escherichia coli*) (green) are shown in the context of bacterial cytoplasmic membrane (gray rectangle) with periplasmic side and cytoplasmic side represented above and below the gray rectangle, respectively. The N- and C-termini of the respective proteins are indicated with “N” or “C”. **(b)** Lysis profiles of assay strain TB28 co-transformed with plasmids carrying inducible *sgl* genes (sgl^M^, sgl^PP7^,and GFP-sgl^PP7^)and compatible plasmids expressing MurJ orthologs (MurJ_TA_ ^28^ and AmJ ^29^). These include pEXT21 + pBAD24 (emplty vector control, red open circle), pEXT21 + pBAD24 *sgl^M^* (dark blue open square), *pEXT21-murJ_TA_* + pBAD24 *sgl^M^* (light orange filled circle), pEXT21 + pBAD24 *sgl^PP7^* (dark green filled square), *pEXT2-murJ_TA_* + pBAD24 *sgl^PP7^* (black cross), pEXT21 + pBAD24 *gfp-sgl^PP7^* (pink triangle), pEXT21-*murJ_TA_* + pBAD24 *gfp-sgl^PP7^* (purple filled circle), pCS83 *amJ* + pBAD24 *sgl^PP7^* (red filled square), pCS83 *amJ* + pBAD24 *gfp-sgl^PP7^* (light blue diamond). **(c)** The amino acid substitutions in *E. coli murJ* (MurJ_EC_) that confer resistance to *gfp-sgl^PP7^* are shown on the crystal structure of an inward open conformation of MurJ_EC_ (PDB 6CC4). The TMDs that line the central hydrophilic cavity are colored as follows TMD1 (olive darb), TMD2 (steel blue), TMD7 (magenta), and TMD8 (gold rod). The substituted amino acid is highlighted as cyan spheres on TMD7 (magenta). Lateral view (left) and periplasmic view (right).

### Suppressor patterns for the L-like Sgl proteins

In contrast to Sgl^M^, and Sgl^PP7^, where MurJ is one of the few genes identified as a suppressor, the suppressor profiles for the presumptive L-like Sgls from MS2, KU1, Hgal1, and PRR1 are rich and complex (Fig 2b, Supplementary Figs. 2-6, Extended Data 5). Aside from *waaQ* and *galE*, ten genes showed high fitness scores indicating they play a role in mitigating the toxicity induced by Sgls: *micF*, *mltF*, *mpl*, *rodZ*, *rbsR*, *treB*, *yajC*, *yceG*, *yqgE* and *ytfP*. *mltF* and *yceG* encode lytic transglycosylases involved in PG turnover^21,30,31^. Mpl is a murein peptide ligase involved in recycling the PG precursors derived from cell wall turnover^32,33^. By providing an alternate source of PG precursors, these three could provide palliative relief indirectly to any developing insult to cell wall biosynthesis and turnover (Fig 1a). MicF is a small RNA and has been shown to regulate OM porin expression and can thus influence the integrity and permeability of the envelope^34^. Other hits are harder to rationalize. YajC is part of the Sec translocon accessory complex, but of unknown function. RbsR is a repressor of ribose catabolism and transport. RodZ is a key regulator of cell division, interacting directly with the FtsZ septal ring. TreB is part of the PTS pathway for trehalose import.None of these genes are essential^35^ and in none of these cases is there a lytic phenotype associated with a gene knockout. The simplest idea is that all of these suppressor fragments exert indirect effects, or act through sRNAs like MicF and RirA (from processing of the waaQ transcript)^34^, many of which likely remain cryptic.

In addition to the broadly high scoring genes mentioned above, we also uncovered high fitness scores for few genes that are specific for some L-like Sgls (such as *hslU, rstA, dacC, slt, sdiA, speA* and *galK*, Fig 2b) and the specific pattern of these scores is hard to rationalize. For example, high scores of *slt* (encoding the main lytic murein transglycosylase) and *dacC* (codes for D-alanyl D-alanine carboxypeptidase) makes sense as they are considered to be part of PG quality-control pathways, but the specific fitness effect of *slt* in the KU1 and MS2 assays compared to that of *dacC* for the Sgls of KU1, Hgal1, and PRR1 dataset is intriguing. It should be noted that the genes encoding a protease active against membrane proteins (for example, HslVU) is over-represented in the suppressor collection for the three out five L-like Sgls but not for Sgl^M^ or Sgl^PP7^ (MS2, Hgal1, and PRR1, Fig 2b). This may indicate that L-like Sgls largely remain sensitive to proteases during the lytic pathway, possibly because they do not form a stable complex with a protein target. Overall, the conclusion is that the lytic function of these four L-like Sgls can be suppressed by multicopies of many genes involved in envelope homeostasis.

## Discussion

In this study we applied an unbiased genome-wide genetic screen to identify multicopy suppressors of Sgl lysis proteins - a diverse group of lysis genes from *Fiersviridae* phages. As a proof-of-principle, we benchmarked our genetic screen against the toxicity of Sgl^M^ and we recapitulated the identification of its known target, the lipid II flippase,MurJ, as the high confidence candidate (Figs. 1c,d) ^19^. Encouraged with this result, we applied the method to Sgl^MS2^, the L protein, whose target has remained enigmatic over half a century. In addition to Sgl^MS2^, we also screened Sgls from four other ssRNA phages: KU1, Hgal1, PRR1 and PP7 that share characteristic 4 motif structure of L protein and therefore were previously proposed to have the same cellular target. In total, we observed 190 high-confidence hits across the *sgl^M^, sgl^PP7^, sgl^PRR1^, sgl^MS2^*, and *sgl^Hgal1^* Dub-seq screens. We followed up with one of the top scoring candidates for Sgl^PP7^ and confirmed that Sgl^PP7^ lethality can be rescued by expression of heterologous lipid II flippase MurJ, strongly suggesting it to be a target of Sgl^PP7^. Thus it appears that, functionally, Sgl^PP7^ is not an L-like Sgl. Though most of the high confidence hits play a role in PG biosynthesis or alleviating outer membrane stress response and validate in follow ups, some of the suppressor hits are difficult to rationalize. Nevertheless, the Dub-seq genetic screen successfully uncovers multicopy suppressors and provides insights into Sgl target/repair pathways at a genome-wide scale that may be challenging to obtain via traditional approaches.

One of the major unexpected findings of this study was that the Sgl^PP7^ targets MurJ, an essential lipid II flippase in Gram-negative bacteria. Interestingly, the Sgl^PP7^ has no primary structure resemblance to the other MurJ-targeting Sgl, Sgl^M^. Furthermore, the lytic function of these two Sgls were blocked by the same single missense change (Q244P) in MurJ, suggesting that these two disparate Sgls have not only convergently evolved to target the same protein but may also target the same molecular interface on MurJ. The resistance allele Q244P is located on TMD7, one of the four TMDs lining the central hydrophilic cavity of MurJ. Interestingly, TMD7 undergoes large conformational changes between periplasmic-open and cytoplasmic-open states of MurJ and Gln244 is positioned at the bend in the helix. Locking MurJ in either of the conformational states leads to accumulation of Lipid II in the inner leaflet of the inner membrane and ultimately results in cell lysis. Previously, cysteine accessibility studies (SCAM) have shown that Sgl^M^ locks MurJ in periplasmic-open conformation and blocks the transfer of Lipid II across the membrane. Given the putative interaction interface of Sgl^PP7^ at the highly dynamic TMD7 of MurJ, potentially Sgl^PP7^ locks MurJ in the opposite conformation to Sgl^M^ i.e, cytoplasmic-open conformation. Future SCAM analysis and structural studies of Sgl^PP7^-MurJ complex should shed light on both the conformation state of MurJ-Sgl^PP7^ complex and its interaction interface.

Recently, the number of experimentally validated Sgls has expanded by 35 and they share no detectable similarity to the previously characterized Sgls ^14^. The high sequence diversity of Sgls naturally implies possible diversity in molecular targets to affect host cell lysis. However, we speculate evolution of Sgls is highly constrained as being part of compact ssRNA phage genomes, in addition to being commonly found as overlapped or encoded within other phage genes. Also, being membrane associated probably makes them target other membrane proteins such as peptidoglycan proteins localized in the inner membrane. Hence, convergent evolution of Sgls to target the limited number of host targets is an inevitable consequence and one should expect more cases of convergent evolution to the known targets such as MurA, MraY, and MurJ. We note here that the recently reported 35 Sgls were selected for having a predicted TMD domain so it is possible they are likely biased towards interacting with MurJ and MraY. The fact that both Sgl^M^ and Sgl^PP7^ target MurJ suggests that there is more than one way to exploit the same “weak spot” in the bacterial cell wall machinery. Furthermore, a target uncovered in one species of bacteria (i.e., MurJ in *Pseudomonas*) could also serve as one in another more distant species (i.e., MurJ in *E. coli*). Thus, by studying convergently evolved Sgls one could gain insights into built-in universal molecular “weak spots’’ across various species.

Here we limited this study to the discovery of suppressors for unique Sgl lysis proteins and *Fiersviridae*. We anticipate this unbiased genetic screening approach to be generalizable and extendable to discover suppressors of many other toxic genes found in nature including Sgls from the recent hyperexpansion of ssRNA phage genomes. Further, this approach could be useful in the study and annotation of dsDNA and ssDNA phage genomes and host-encoded small toxic genes ^36^. We demonstrate here that by repurposing Dub-seq technology for carrying out unbiased and high-throughput suppressor screens will greatly expedite hypothesis generation and target identification of Sgl lysis proteins, providing a new avenue for antibiotic and phage-derived biotechnological discovery.

## Author contributions

B.A.A. conceived the project.

B.A.A., H.C., and V.K.M built and characterized the Dub-seq libraries used in this study.

B.A.A., K.C., H.C., and J.K., designed constructs, performed experiments, processed, and analyzed data.

A.M.D. provided critical reagents, support for sequencing efforts, and advice.

B.A.A., K.C., R.F.Y., V.K.M., and A.P.A. wrote the paper.

V.K.M., and A.P.A. supervised the project

## Acknowledgments

The authors gratefully thank Simon Roux (Joint Genome Institute) for helpful discussions at various stages of this project. This project was funded by the Microbiology Program of the Innovative Genomics Institute, Berkeley. The initial Dub-seq library characterization for this project was funded by ENIGMA, a Scientific Focus Area Program at Lawrence Berkeley National Laboratory, supported by the U.S. Department of Energy, Office of Science, Office of Biological and Environmental Research under contract DE-AC02-05CH11231. Sequencing was performed at: Vincent J. Coates Genomics Sequencing Laboratory (University of California at Berkeley), supported by NIH S10 Instrumentation Grants S10RR029668, S10RR027303, and OD018174. R.F.Y acknowledges funding from NIGMS grant R35GM136396.

## Methods

### Bacterial strains and growth conditions

In general, all *E. coli* strains were grown at 37 °C, 180 rpm in Lysogeny Broth (LB-Lennox broth, Sigma) supplemented with antibiotics, unless stated otherwise. When appropriate, 50 μg/mL kanamycin and/or 34 μg/mL chloramphenicol (denoted with +K or +C, respectively) were added to media. All bacterial strains and libraries were stored at - 80°C for long term storage in 25% sterile glycerol (Sigma). All library assays were performed in NEB10beta strain backgrounds (*araD139* Δ(*ara-leu*)7697 *fhuA lacX74 galK* (Φ80 Δ(*lacZ)M15*) *mcrA galU recA1 endA1 nupG rpsL* Δ(*mrr-hsdRMS-mcrBC*), New England Biolabs). For a complete list of strains and plasmids, please refer to Extended Data 6.

### Construction of sgl expression strains

Template sequences for *sgl^Hgal1^, sgl^M^, sgl^MS2^, sgl^PRR1^* and *sgl^PP7^were* identified from the NCBI-deposited genomes: NC_019922, NC_019707, NC_001417, NC_008294, and NC_001628, respectively. As a toxic gene control, we used protein PC02664 detected from a phage genome infecting *E. coli* ^37,38^. Each gene was codon-optimized for *E. coli*, had BsaI sites removed, and synthesized *de novo* (IDT, TWIST Bioscience). *Sgl* genes were cloned into pBA368, a golden gate *gfp*-dropout vector derived from pBbA2K-rfp. DNA assembly was performed via golden gate assembly using BsaI (New England Biolabs), pBA368, and one of the synthesized *sgls*. Reactions were cleaned up using Clean and Concentrate (Zymo), transformed into NEB10beta competent cells (New England Biolabs), and plated on LB+K. GFP-colonies were picked, grown up, stored at −80°C, and verified for the intact *sgl*.

For all strains lytic activity was measured via plate reader assay before constructing suppressor libraries. Strains were inoculated into LB+K media overnight. Cells were diluted 50X into LB+K media with varying levels of anhydrotetracycline (aTc) ranging from 0-200ng/mL in a flat bottom 96-well plate (Corning 3904). Sgl-mediated lysis progressed in Tecan Infinite F200 readers with orbital shaking and OD600 readings every 15 min for 3-5 hours at 37°C. Strains with functional Sgl phenotypes typically had visible lysis after ~90 minutes.

Plasmid pBAD24-*sgl^PP7^-lacZ_⍰_* was constructed in multiple steps. First, the *sgl^PP7^* (NC_001628.1) was codon-optimized for *E. coli* expression (IDT) and a synthetic DNA construct was obtained (GenScript). The synthetic *sgl^PP7^* DNA was amplified using primers KC94 and KC116 and the resulting PCR product was gel purified (Qiagen), digested with restriction enzymes EcoRI and XhoI (New England Biolabs), and sub-cloned into plasmid pKC3, replacing *sgl^M^* in pKC3.

Plasmid *pBAD24-gfp-sgl^PP7^-lacZ_⍰_* was constructed via the Overlap Extension PCR method using primers KC36 and KC127 to amplify a *gfp* megaprimer in the first PCR ^39^. The megaprimer was then used to insert *gfp* into *pBAD24-sgl^PP7^-lacZ_⍰_* plasmid during the second PCR. The product of the second PCR reaction was treated with DpnI and then transformed into competent XL1Blue cells. The constructs were verified by sequencing (Eton Biosciences) with primers KC30 and KC31.

### Construction of Dub-seq suppressor libraries

Dub-seq suppressor libraries were constructed by transforming the plasmid Dub-seq library, pFAB5516 ^15^, directly into *sgl* expression strains (above) via electroporation. Competent cells were created from an overnight culture diluted 70X into 25mL LB+K and shook at 37°C, 180rpm for ~3 hours until OD600 0.5-0.7. The resulting mid-log cultures were chilled at 4°C. Cultures were centrifuged (Beckman-Coulter Allegra 25R) for 5 minutes at 8000xg and subjected to three washes: (i) once with 25 ml chilled water (ii) and twice with 15 ml chilled 10% glycerol. The cell pellets after the final glycerol wash were resuspended in 10% glycerol, yielding ~250 ul of cells.

For each *sgl* library, 5 parallel transformations were performed to minimize inefficiency bias from any individual transformation. Each transformation consisted of 40μL of competent cells and 10ng of the pFAB5516 plasmid library transferred to a chilled cuvette (1mm gap, VWR). Cuvettes were electroporated using a BTX™-Harvard Apparatus ECM™ 630 Exponential Decay Wave Electroporator with the following parameters: voltage (1800 V), resistance (200 Ω), and capacitance (25 μF). Following each transformation, cells were recovered in 1mL LB+K media at 37°C for 1 hour. For each transformation, 980μL of each recovery was plated and spread out onto LB+K+C Agar in a 245mm x 245mm bioassay dish (Nunc). The remaining 20μL of cells were serially diluted and plated onto a standard LB+K+C Agar plate to estimate the number of transformants per electroporation. All transformations were incubated at 37°C overnight.

After overnight incubation at 37°C, we first quantified the transformations to ensure we had at least 250,000 total estimated colonies (i.e., ≥5X pFAB5516 library coverage). We then picked 10 colonies from each of the transformations and PCR, followed by Sanger sequencing to ensure that the *sgl* was free of mutations. If any *sgl* mutations were detected in this subset, we repeated library construction. The transformant colonies were scraped and resuspended in 25mL LB+K+C media and processed as described above to make multiple 1mL −80°C freezer stocks ^15^. Because pFAB5516 was characterized earlier ^15^, there was no need to perform library mapping PCRs at this step. An overview of library composition is summarized in Supplementary Table 1 and a gene-level description is shown in Extended Data 2.

### Liquid culture fitness experiments

Competitive fitness experiments were performed in liquid culture with two replicate experiments were performed per *sgl* suppressor experiment. Briefly, a 1 mL aliquot of suppressor Dub-seq library was gently thawed and used to inoculate a 25 mL of LB+K+C media. The library culture was grown to an OD600 of ~1.0 at 37°C. From this culture two 1 mL pellets were collected, comprising the ‘Time-0’ or reference samples in BarSeq analysis. The remaining cells were diluted to a starting OD600 of 0.02 in LB+K+C media. A 690 μL volume of cells was mixed with 10μL of diluted aTc (Sigma) and transferred to a 48-well microplate (700 μL per well) (Greiner Bio-One #677102) and covered with breathable film (Breathe-Easy). For all experiments, unless otherwise noted, aTc was used at 15.6 ng/mL. The progress of Sgl lysis was followed in Tecan Infinite F200 readers with orbital shaking and OD600 readings every 15 min for 8-12 hours at 37°C. At the end of the experiment, contents of each well were collected and spun down on a table top centrifuge to collect as a pellet individually. All pellets were stored at −80°C until prepared for BarSeq (detailed below). A summary of all library experiments is described in Extended Data 1.

### BarSeq of Dub-seq pooled fitness assay samples

Plasmid DNA was isolated from stored pellets of enriched and ‘Time 0’ (“time=zero”) Dub-seq samples using the QIAprep Spin Miniprep Kit (Qiagen). We performed 98°C BarSeq PCR protocol as described previously ^40^. BarSeq PCR in a 50 μL total volume consisted of 20 μmol of each primer and 150 to 200 ng of plasmid DNA. For the HiSeq4000 runs, we used an equimolar mixture of four common P1 oligos for BarSeq, with variable lengths of random bases at the start of the sequencing reactions (2–5 nucleotides). Equal volumes (5 μL) of the individual BarSeq PCRs were pooled, and 50 μL of the pooled PCR product was purified with the DNA Clean and Concentrator kit (Zymo Research). The final BarSeq library was eluted in 40 μL water. The BarSeq samples were sequenced on Illumina HiSeq4000 with 50 SE runs.

### Data processing and analysis of BarSeq reads

Fitness data for Dub-seq suppressor libraries were analyzed as previously described with a few modifications as described, using *barseq* script from the *Dub-seq* python library with default settings ^15^. From a reference list of barcodes mapped to the genomic regions (BPSeq and BAGseq), and the barcode counts in each sample (BarSeq), we estimated fitness values for each genomic fragment using the *gscore* script from the Dub-seq python library. At this step, instead of pooling all Time0 samples together, the time-zero samples within each suppressor library were pooled, since the composition and abundance of library members between libraries was distinct. For instance, MS2 experiments had time-zero samples different from those of the PP7 experiments. The *gscore* script identifies a subset of barcodes mapped to the genomic regions that are well represented in the time-zero samples for a given experiment set. A barcode was required to have at least 10 reads in at least one time-zero (sample before the experiment) sample to be considered a valid barcode for a given experiment set. The *gscore* script was used to calculate a fitness score (normalized ratio of counts between the treatment sample and sum of counts across all time-zero samples) for the strains with valid barcodes. From the fitness scores calculated for all Dub-seq fragments, a fitness score for each individual gene that is covered by at least one fragment was calculated using nonnegative least squares regression ^15^. The nonnegative regression determines if the high fitness of the fragments covering the gene is due to that particular gene or its nearby neighboring gene, and avoids overfitting. Raw data for reads, fscores, and gscores across all experiments are provided in Extended Data 3, Extended Data 4, and Extended Data 5, respectively.

We applied additional filters to ensure that the fragments covering the gene had a genuine benefit. Briefly, we identified a subset of the effects to be reliable if the fitness effect was large relative to the variation between start samples (|score| ≥ 2) for both mean and gene fitness scores, the gscores and fscores appeared to be reproducible across replicate experiments, and the number of reads for those fragments was consistently sufficient for the gene score to have little noise. Due to the strong selection pressure and subsequent fitness distribution skew resulting from Sgl activity, all candidate genes passing these filters were then subjected to manual scrutiny. For each gene, all barcodes were analyzed by fscore and reads. Several genes covered by few fragments (ie ≤ 3), had inconsistent fscores, with orders of magnitude different read depth. This bias yielded inflated gscores and were discarded from further analysis. However, genes covered by individual fragments were kept in such cases.

### ASKA-based validations

To validate select lysis suppressor phenotypes from suppressor screens we performed plate reader assays using additional plasmids derived from the overexpression ASKA library ^25^. ASKA plasmids were miniprepped from the ASKA collection using a QIAprep Miniprep kit (Qiagen), transformed into the corresponding *sgl* expression strain, and plated on LB+K+C Agar. Transformants were verified by Sanger sequencing.

Plate reader assays for validations were performed as follows. Strains were inoculated into LB+K+C overnight. Cells were diluted 50X into LB+K+C media and allowed to grow at 37°C, 180rpm to OD600 = 0.5. Cells were then transferred to a Corning 3904 96-well plate (Corning) and induced with varying levels of atc ranging from 0-250ng/mL for *sgl* expression and varying levels of IPTG ranging from 0-200μM for ASKA gene expression. Sgl lysis progressed in Tecan Infinite F200 readers with orbital shaking and OD600 readings every 10 min for 3-5 hours at 37°C. Strains with unsuppressed lysis phenotypes typically had visible lysis after ~90 minutes.

### Suppression by heterologous flippase genes

Strain TB28 was co-transformed with plasmids carrying inducible *sgl* genes (*sgl^M^, sgl^PP7^*, and *GFP-sgl^PP7^*) and compatible plasmids expressing MurJ orthologs (MurJ_TA_ ^28^ and AmJ ^29^) and selected on LB-Amp-Spec-IPTG (100μM) agar plates. The transformants were grown overnight at 37□°C with the same selective media and on the following day 1:200 dilutions of the overnights were added to 25 mL LB with appropriate antibiotics and IPTG (100μM) in a 250 mL flask and grown at 37□°C in an orbital shaker (New Brunswick Co gyrotory water bath shaker model G76) at 250 rpm. The cultures were induced with 0.4% w/v L-arabinose (Sigma-Aldrich) at OD550 ~0.2. The growth was plotted using Kaleidagraph 4.03 (Synergy Software).

### Isolation of GFP-Sgl^PP7^-resistant mutants

Cultures of XL1-Blue *pBAD24-gfp-sgl^PP7^-lacZ_⍰_* were grown overnight at 37□°C with aeration. To perform the Sgl screen/selection, 100□μl of overnight culture was mixed with 400□μl LB and plated on LB-Ara-Amp-IPTG-X-gal agar plates (100 mm). After overnight incubation at 37□°C, blue colonies were picked and purified on the same selection media. The Sgl^PP7^-resistant colonies were grown overnight and both genomic (Qiagen QIAamp DNA micro kit) and plasmid (Qiagen mini prep kit) DNA were extracted. To rule out possible mutations in the lysis gene the plasmid was sequenced with primers KC30 and KC31. The *murJ* locus in the gDNA of the Sgl^PP7^-resistant mutants was amplified by PCR using Phusion high-fidelity DNA polymerase (New England Biolabs) with the primers KC230 and KC234. The amplified PCR product was gel-purified and sequenced with the primers KC230, KC231, KC232, KC233, and KC234.

## Data availability

Sequencing data have been uploaded to the Sequence Read Archive under BioProject accession number PRJNA800467 [http://www.ncbi.nlm.nih.gov/bioproject/800467].

