## Supplementary Information for "Parallel multicopy-suppressor screens reveal convergent evolution of phage-encoded single gene lysis proteins"

*Correspoding authors

 (VKM);

 (APA)

**Contents:**

**Supplementary Table 1: Overview of Dub-seq library composition for all libraries employed in this study.**

**Supplementary Fig. 1: Detailed Dub-seq suppressor screening results against Sgl^M^.**

**Supplementary Fig. 2: Detailed Dub-seq suppressor screening results against Sgl^Hgal1^.**

**Supplementary Fig. 3: Detailed Dub-seq suppressor screening results against Sgl^Ku1^.**

**Supplementary Fig. 4: Detailed Dub-seq suppressor screening results against Sgl^MS2^.**

**Supplementary Fig. 5: Detailed Dub-seq suppressor screening results against Sgl^PP7^.**

**Supplementary Fig. 6: Detailed Dub-seq suppressor screening results against Sgl^PRR1^.**

**Supplementary Fig. 7: Sequence alignment of *Fiersviridae* lysis proteins.**

***Supplementary Table 1. DubSeq suppressor libraries employed in this study.***

The source genome for all libraries is BW25113 with 4509 potential genes for inclusion in Dub-seq experiments. The original pFAB5516 Dub-seq library is included as a reference.

| Library | Sgl Name? ^[1]^ | Established Sgl Target? | Source Phage Genome | Number Fragments | Median Fragment Length (bp) | Genes Covered? | Reference |
| --- | --- | --- | --- | --- | --- | --- | --- |
| pFAB5516 | N/A | N/A | N/A | 27778 | 2514 | 4336 | Mutalik et al., 2019 |
| *sgl^Hgal1^* + pFAB5516 | Sgl^Hgal1^ | No | NC_019922 | 15639 | 2439 | 3736 | This study |
| *sgl^Ku1^* +  pFAB5516 | Sgl^Ku1^ | No | AF227250.1 | 12916 | 2426 | 4042 | This study |
| *sgl^M^* + pFAB5516 | LysM, Sgl^M^ | MurJ | NC_019707 | 17007 | 2455 | 3846 | Chamakura et al., 2017, This study |
| *sgl^MS2^* + pFAB5516 | L, Sgl^MS2^ | No | NC_001417 | 16208 | 2443 | 3802 | This study |
| *sgl^PRR1^* + pFAB5516 | Sgl^PRR1^ | No | NC_008294 | 18477 | 2453 | 3924 | This study |
| *sgl^PP7^* + pFAB5516 | Sgl^PP7^ | No | NC_001628 | 20940 | 2463 | 3968 | This study |

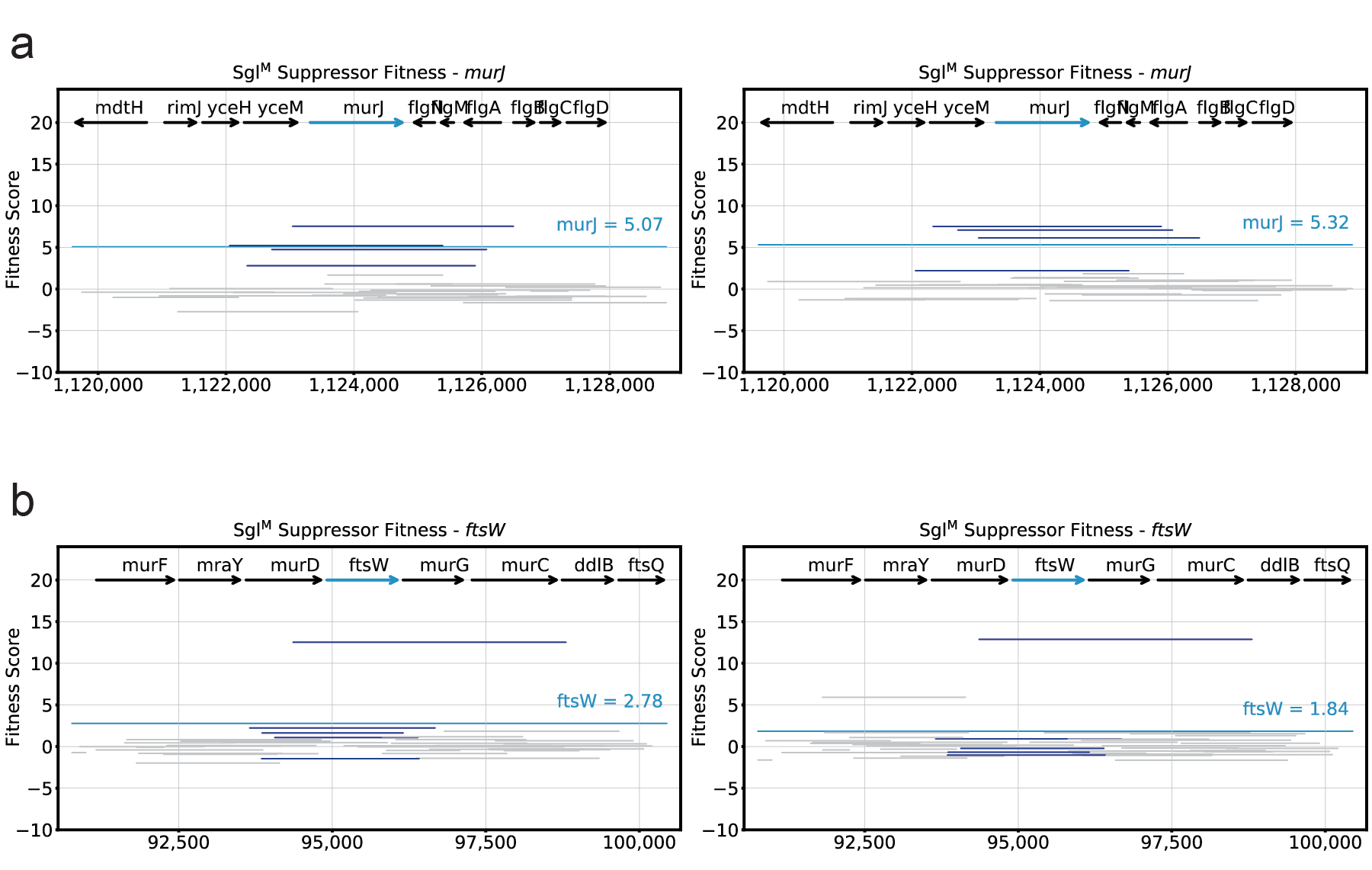

**Supplementary Fig. 1: Detailed Dub-seq suppressor screening results against Sgl^M^.**

(a-b) Dub-seq plots for suppressor screening against Sgl^M^ zoomed in on a gene of interest with experimental replicates shown side-by-side. Genes of interest are significant hits shown in Fig. 2 or candidate multi-gene multi-copy suppressors. Black arrows represent tracked ORFs during analysis. Teal arrows represent the gene of interest. Dark blue lines represent Dub-seq fragments covering the gene of interest with scores shown as *fscores*. Gray lines represent Dub-seq fragments that do not entirely cover the gene of interest with scores shown as *fscores*. Teal lines represent the gene *gscore* of the gene of interest.

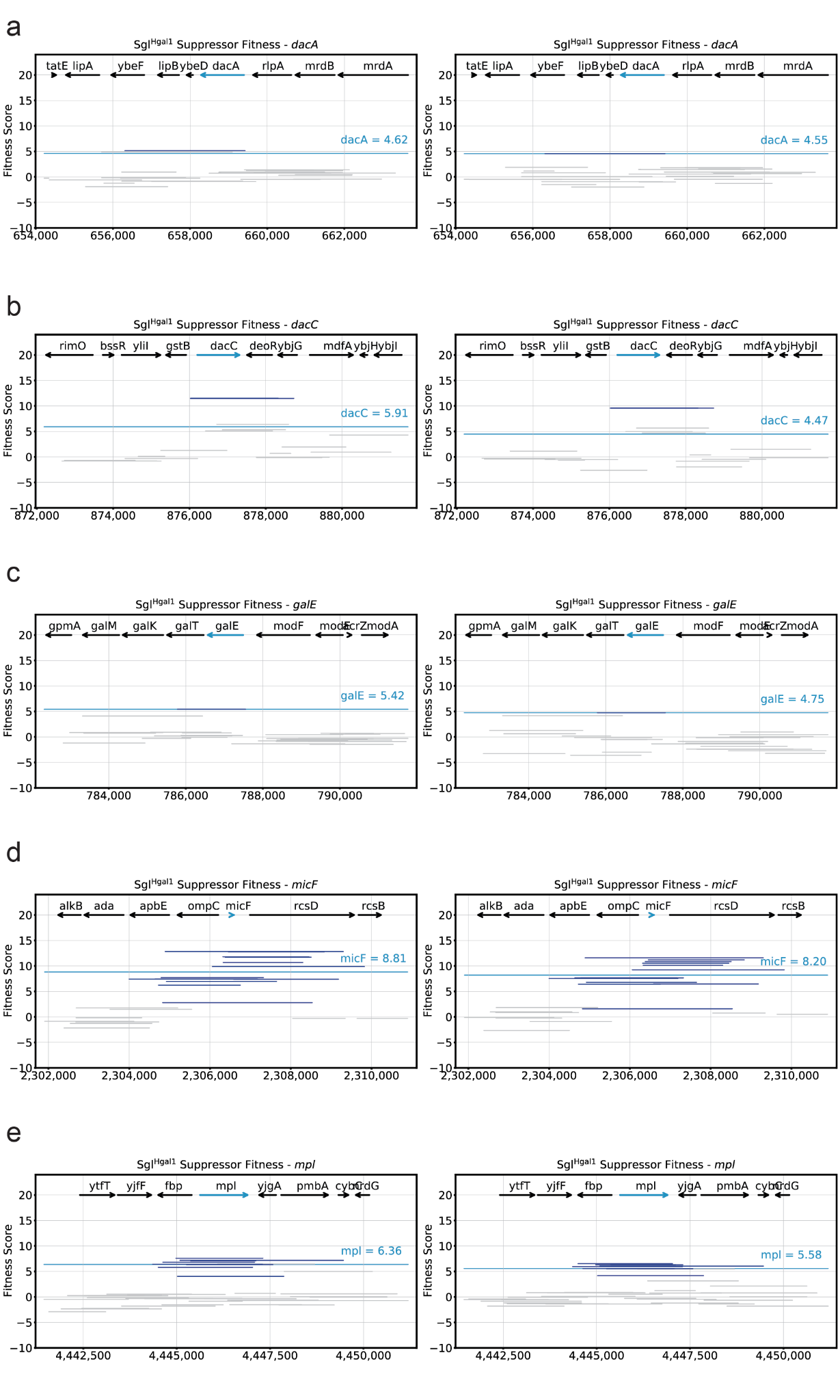

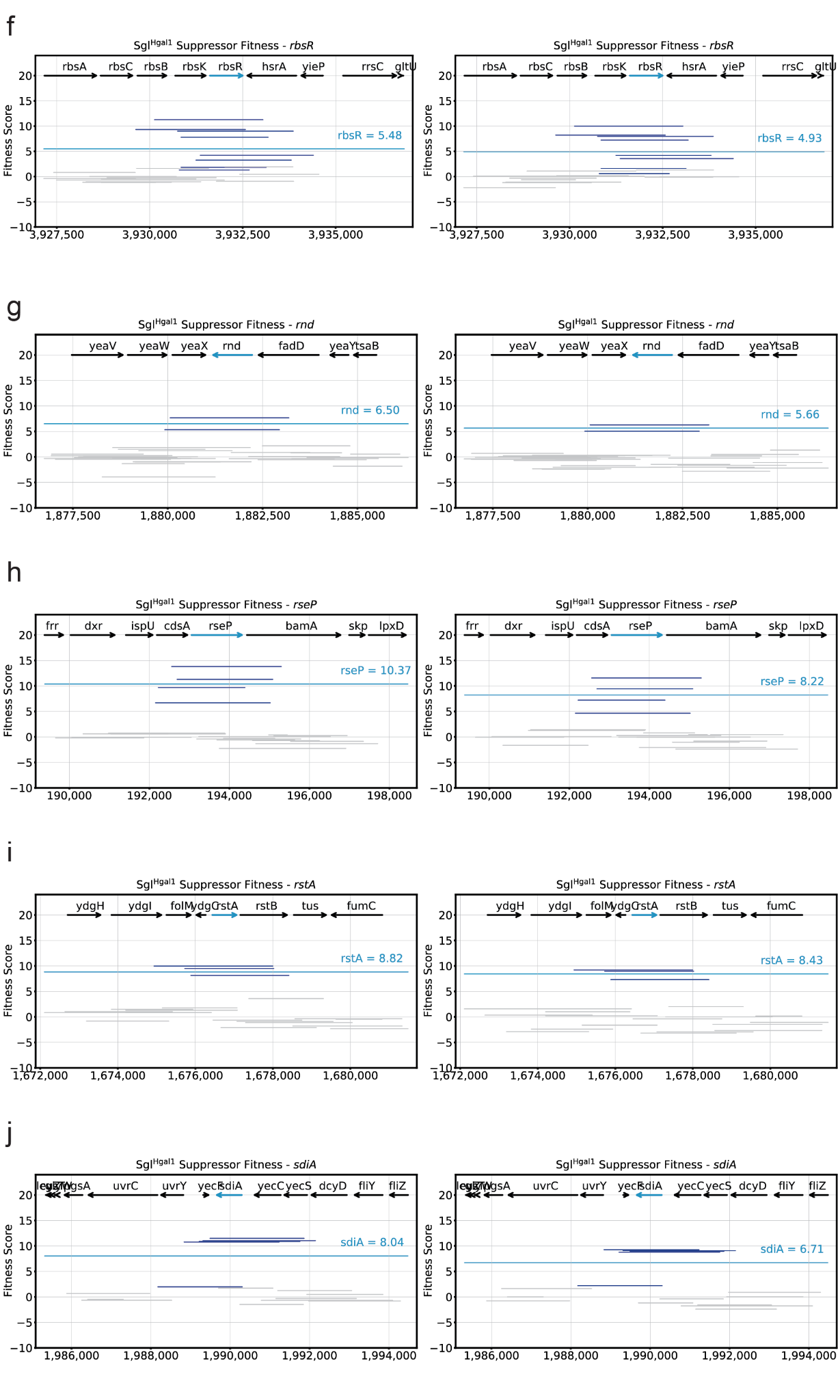

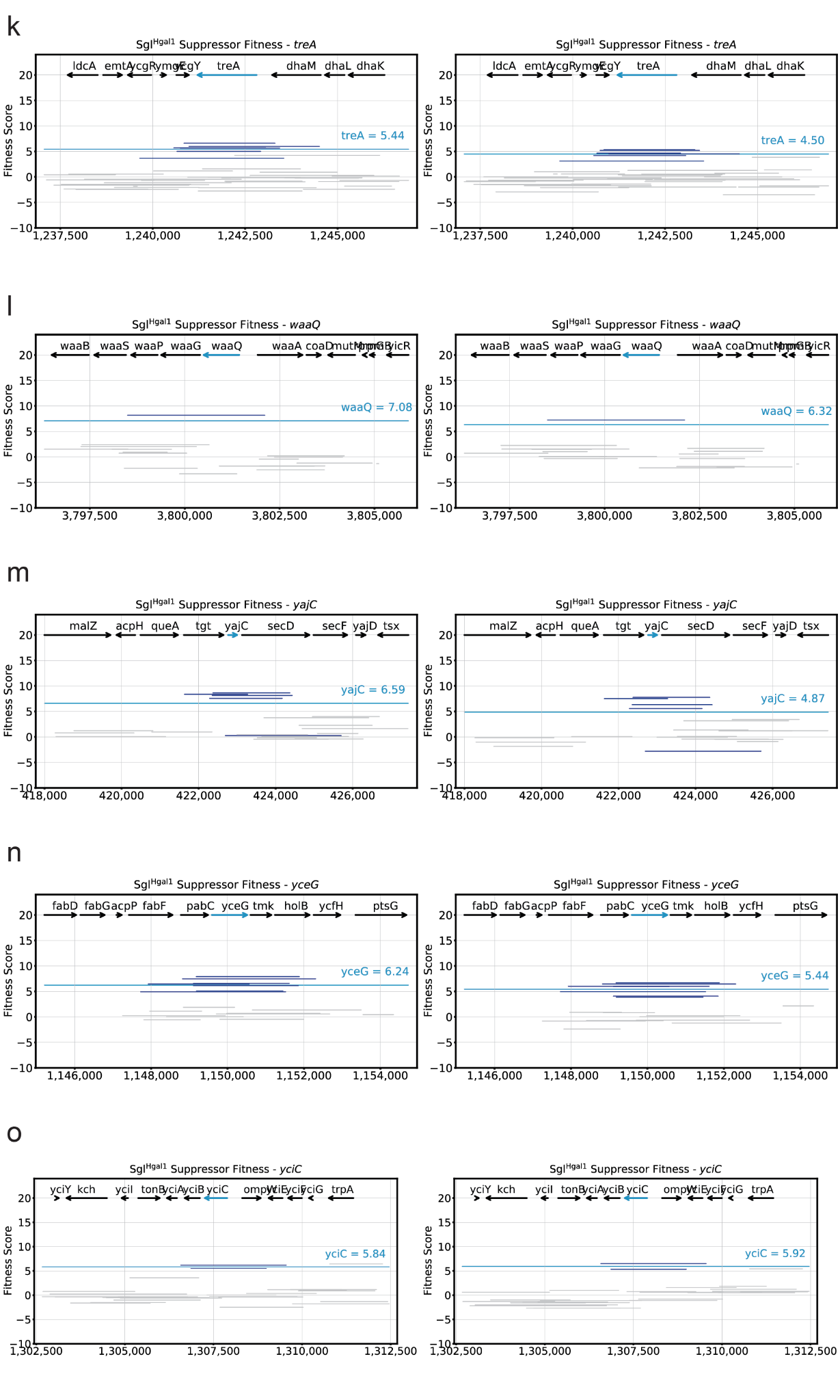

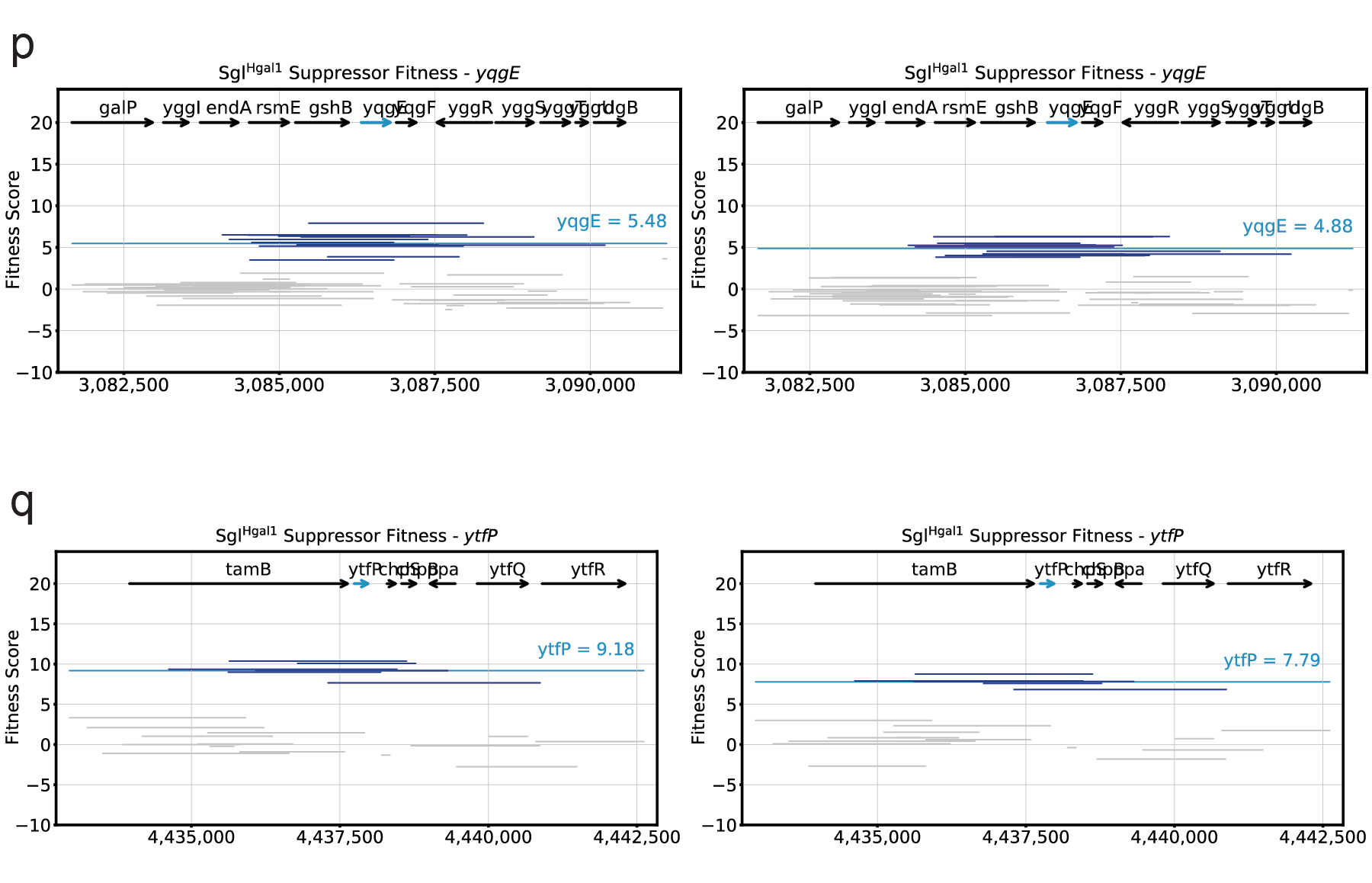

**Supplementary Fig. 2: Detailed Dub-seq suppressor screening results against Sgl^Hgal1^.**

(a-q) Dub-seq plots for suppressor screening against Sgl^Hgal1^ zoomed in on a gene of interest with experimental replicates shown side-by-side. Genes of interest are significant hits shown in Fig. 2 as candidate multi-gene multi-copy suppressors. Black arrows represent tracked ORFs during analysis. Teal arrows represent the gene of interest. Dark blue lines represent Dub-seq fragments covering the gene of interest with scores shown as *fscores*. Gray lines represent Dub-seq fragments that do not entirely cover the gene of interest with scores shown as *fscores*. Teal lines represent the *gscore* of the gene of interest.

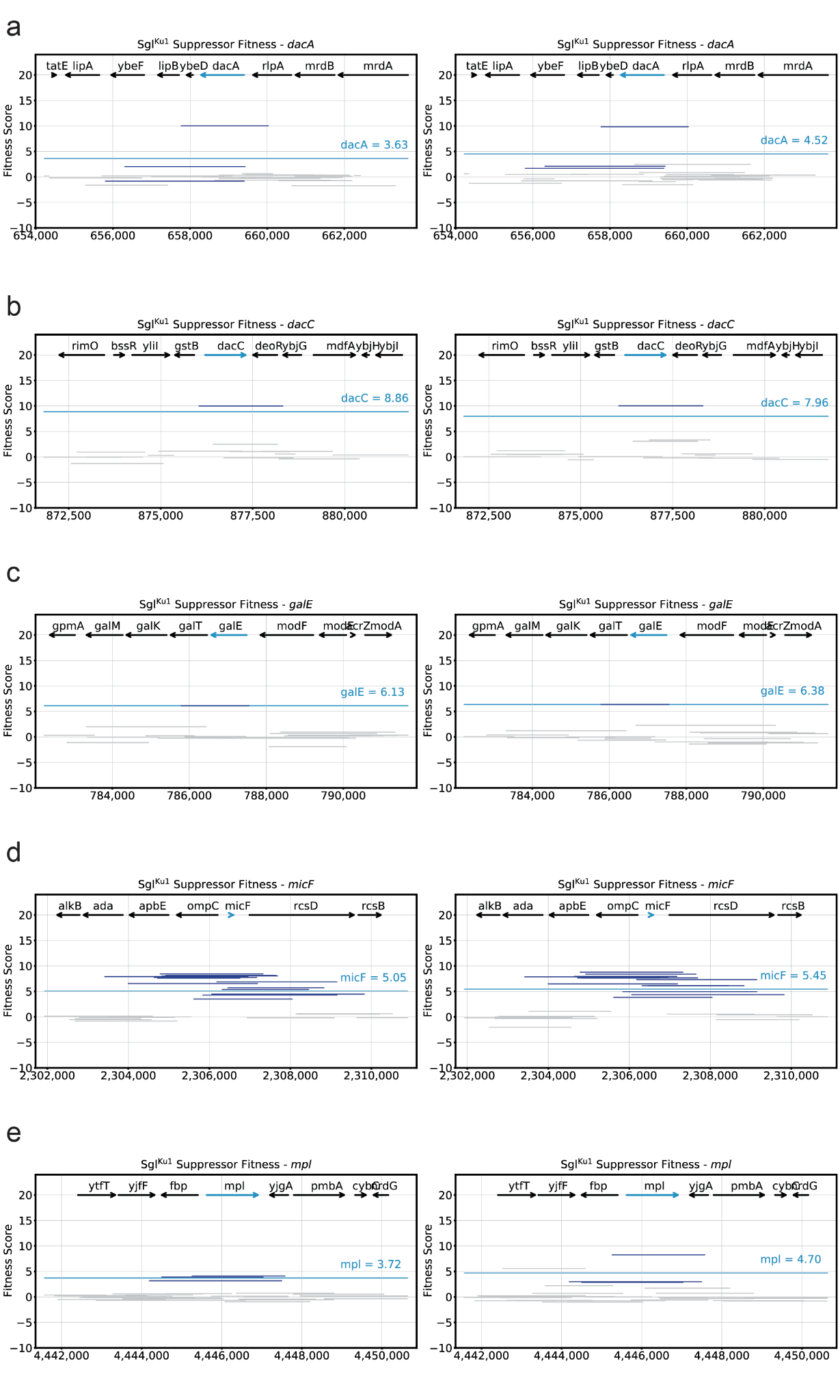

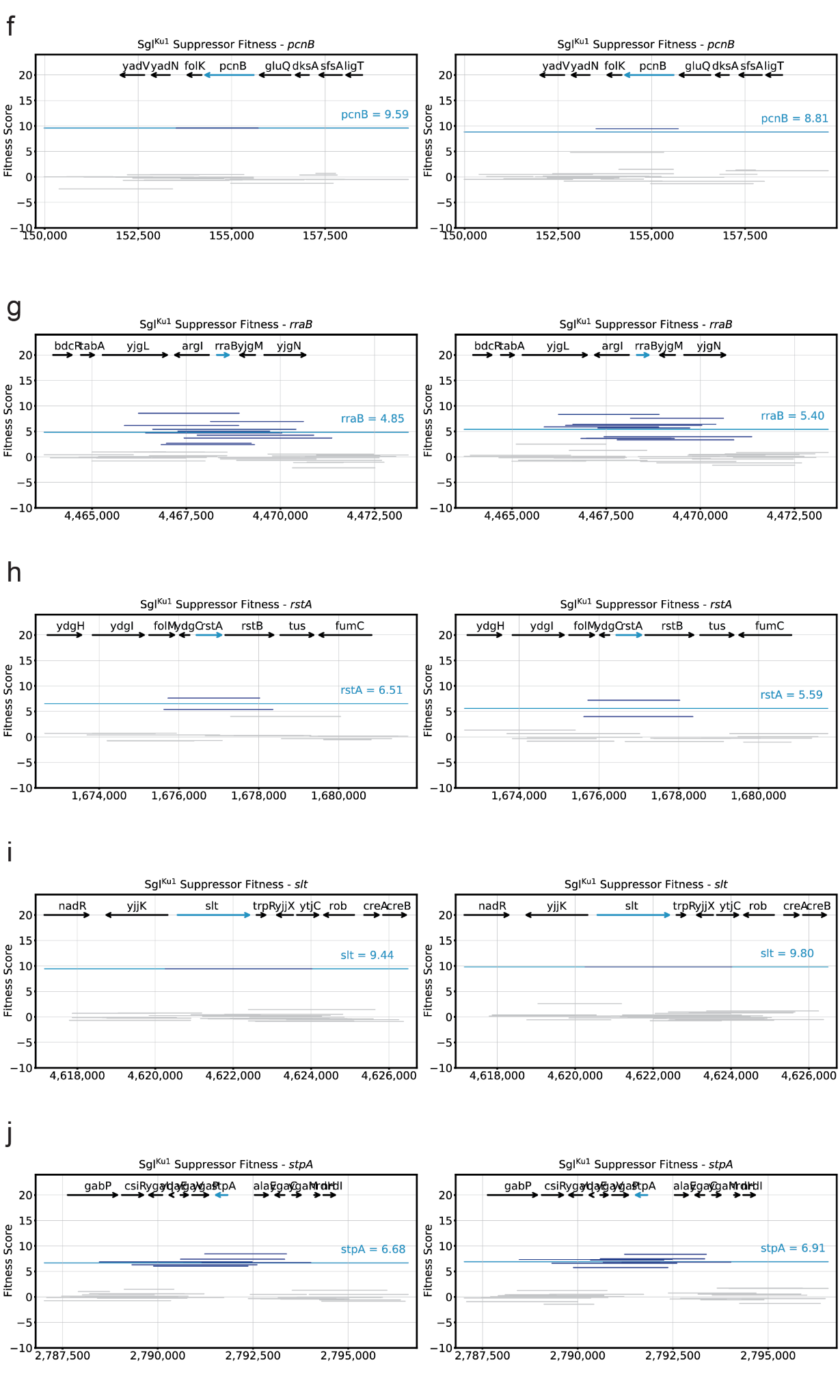

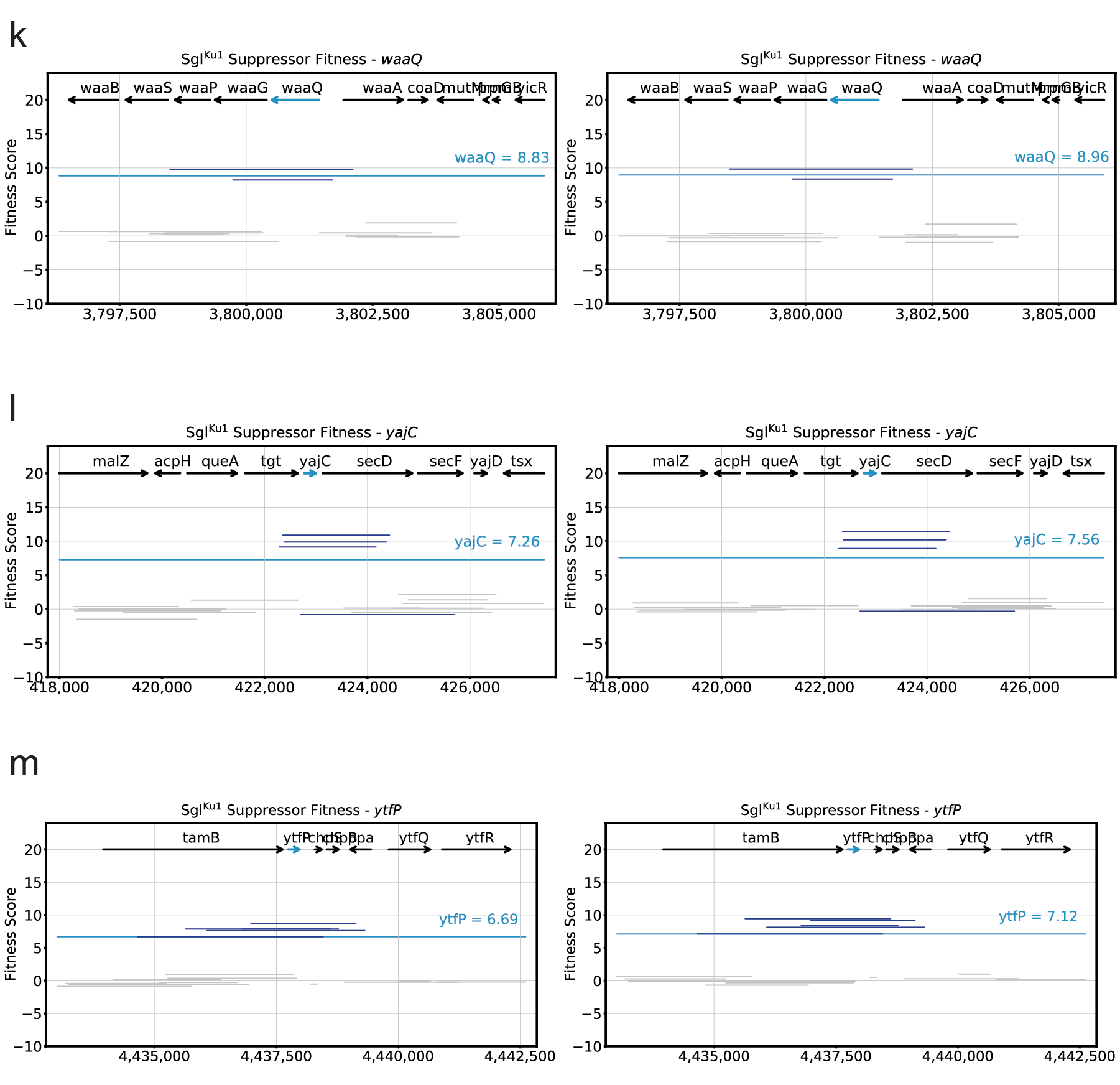

**Supplementary Fig. 3: Detailed Dub-seq suppressor screening results against Sgl^Ku1^.**

(a-m) Dub-seq plots for suppressor screening against Sgl^Ku1^ zoomed in on a gene of interest with experimental replicates shown side-by-side. Genes of interest are significant hits shown in Fig. 2 or candidate multi-gene multi-copy suppressors. Black arrows represent tracked ORFs during analysis. Teal arrows represent the gene of interest. Dark blue lines represent Dub-seq fragments covering the gene of interest with scores shown as *fscores*. Gray lines represent Dub-seq fragments that do not entirely cover the gene of interest with scores shown as *fscores*. Teal lines represent the *gscore* of the gene of interest.

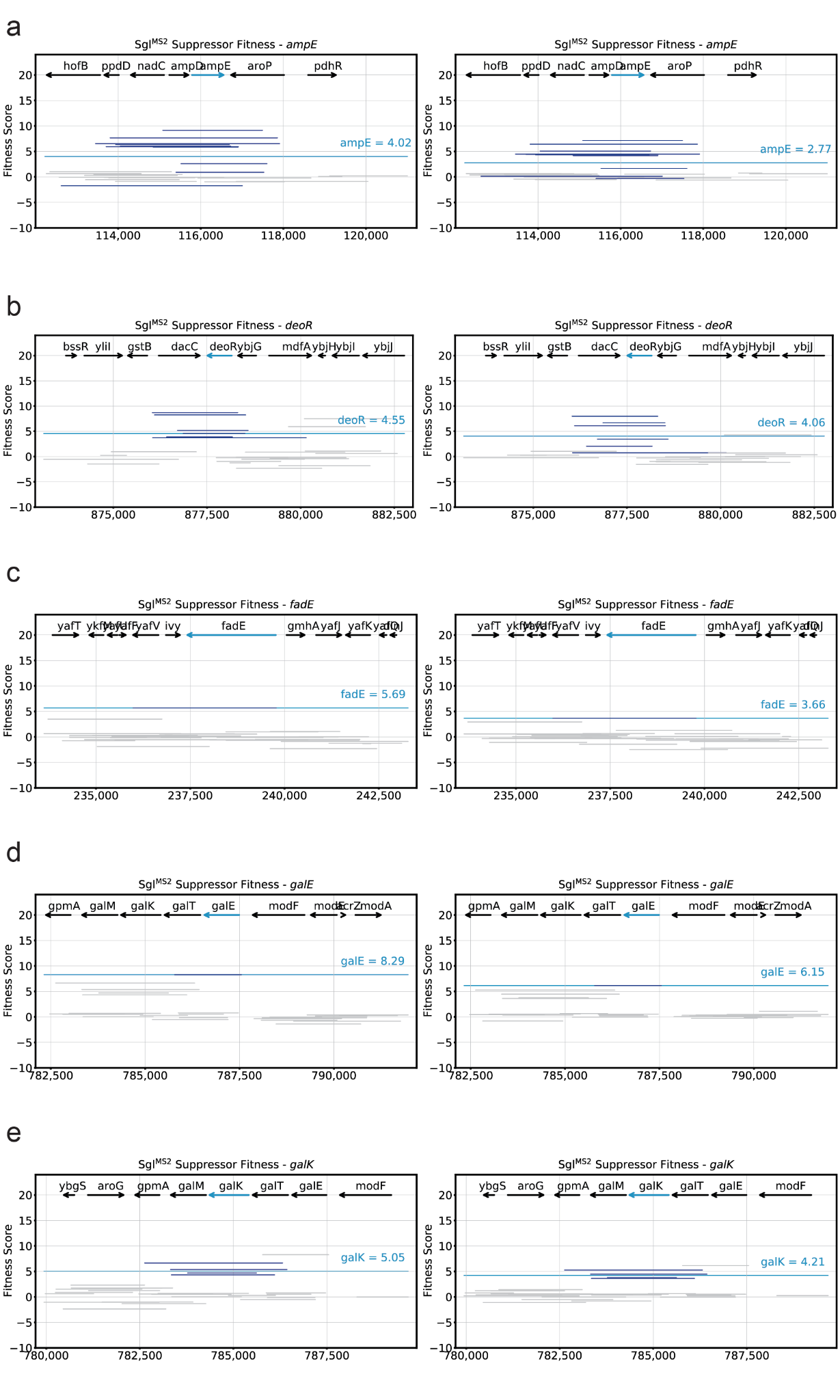

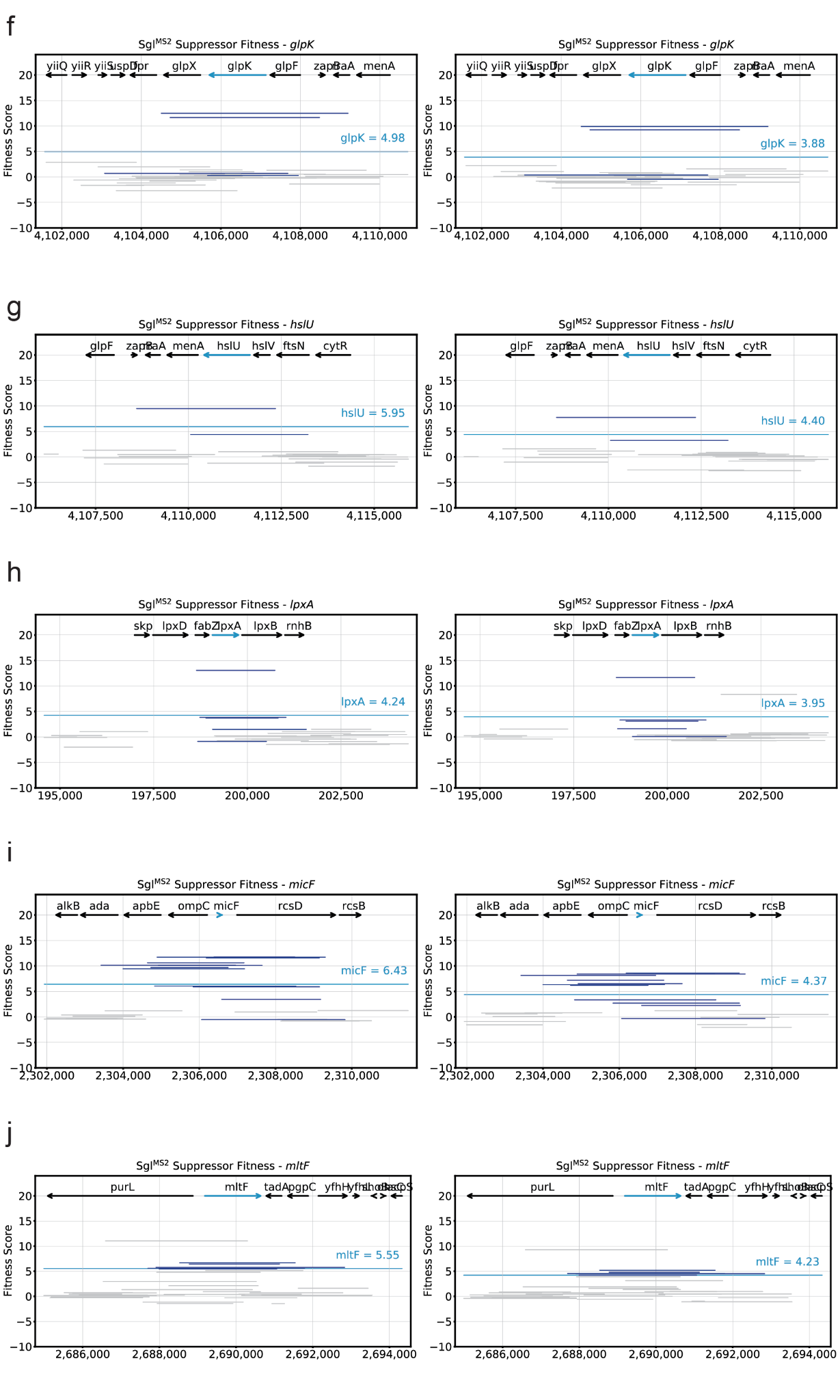

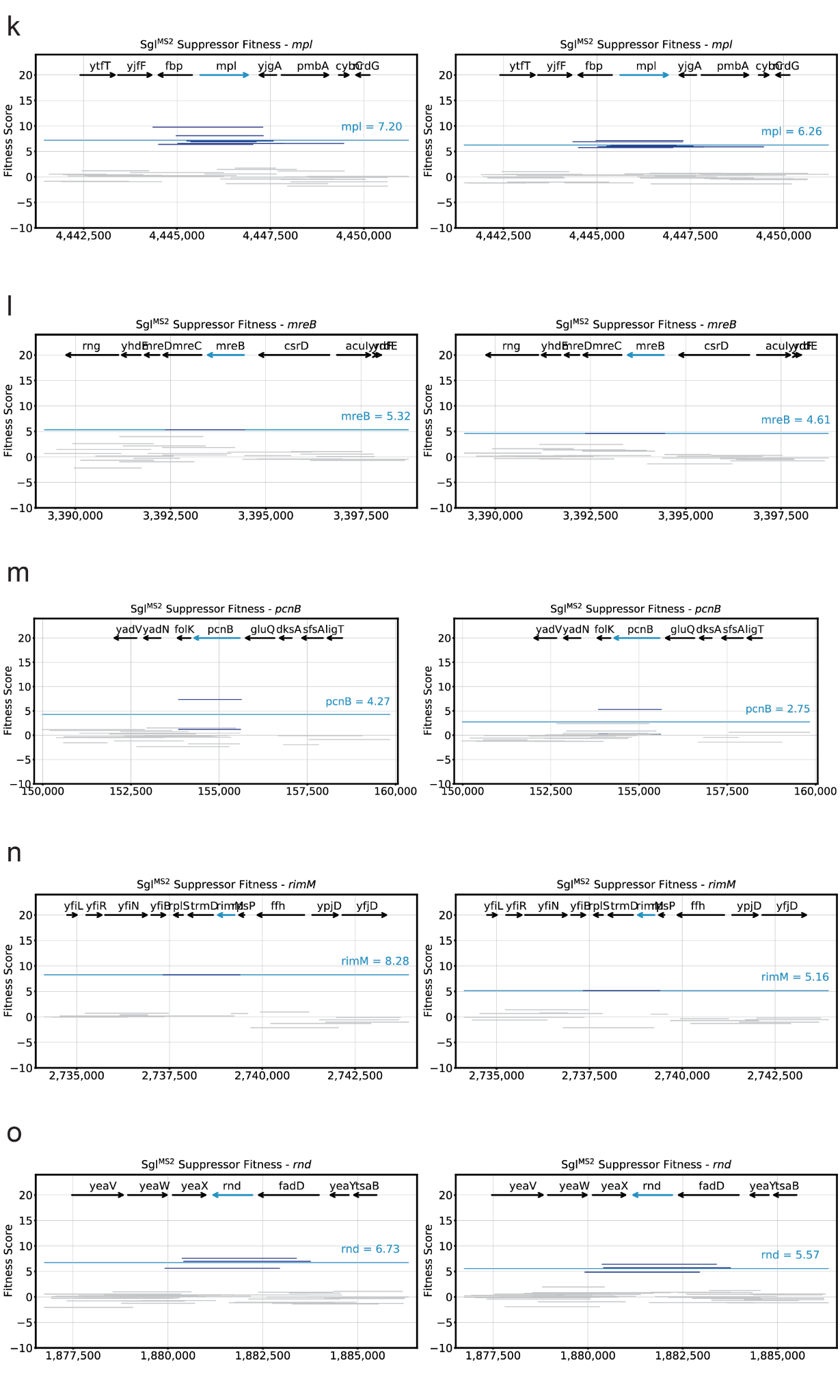

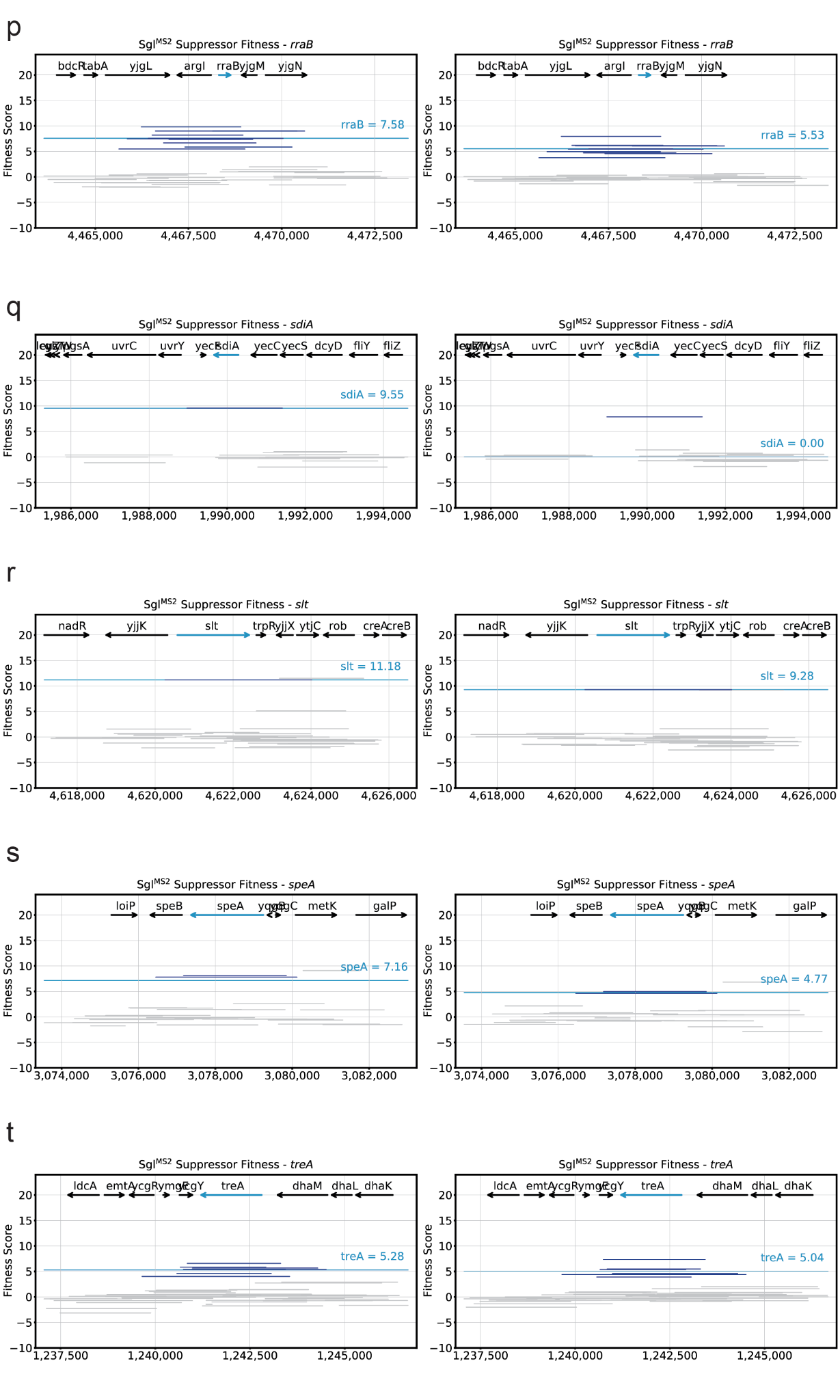

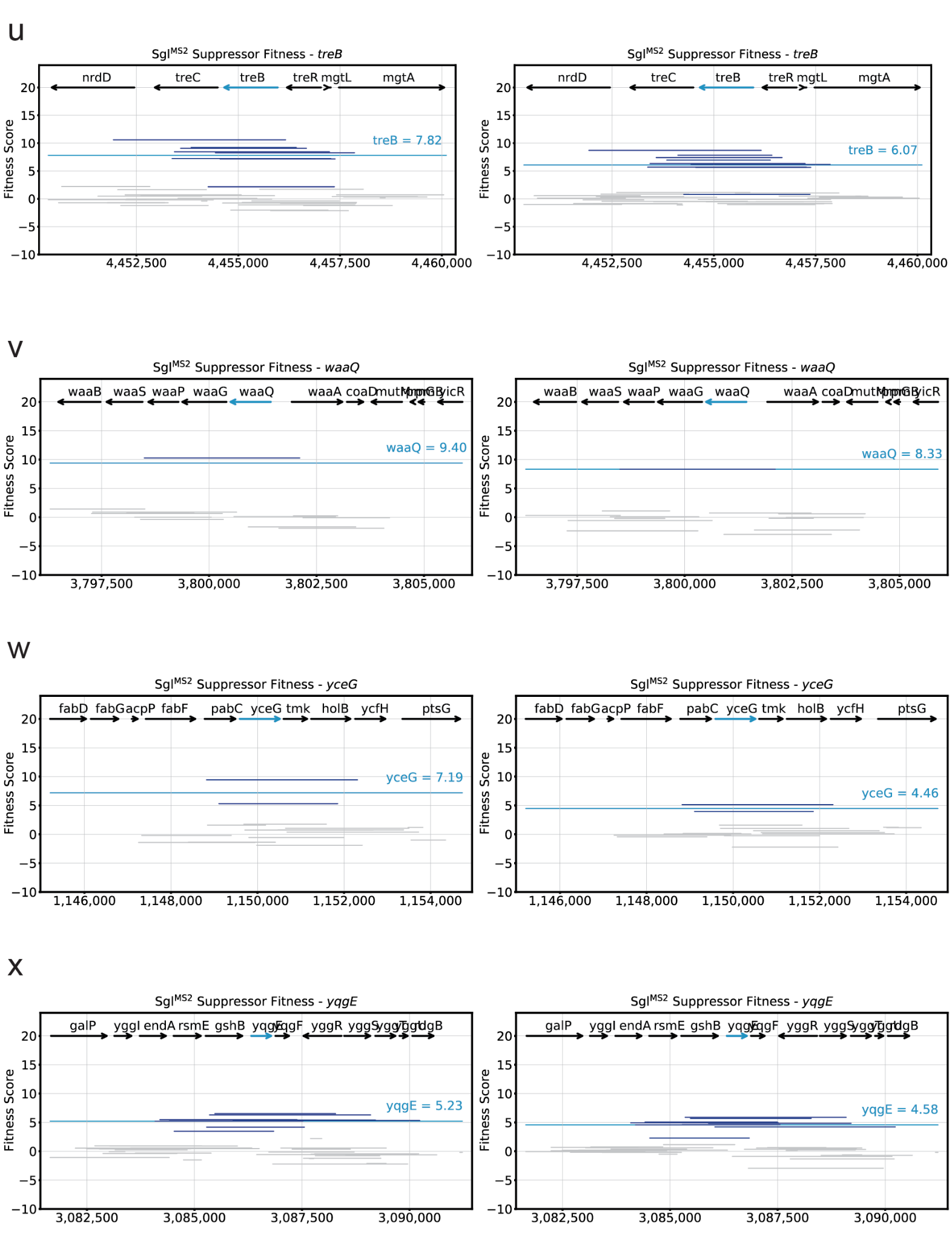

**Supplementary Fig. 4: Detailed Dub-seq suppressor screening results against Sgl^MS2^.**

(a-x) Dub-seq plots for suppressor screening against Sgl^MS2^ zoomed in on a gene of interest with experimental replicates shown side-by-side. Genes of interest are significant hits shown in Fig. 2 or candidate multi-gene multi-copy suppressors. Black arrows represent tracked ORFs during analysis. Teal arrows represent the gene of interest. Dark blue lines represent Dub-seq fragments covering the gene of interest with scores shown as *fscores*. Gray lines represent Dub-seq fragments that do not entirely cover the gene of interest with scores shown as *fscores*. Teal lines represent the *gscore* of the gene of interest.

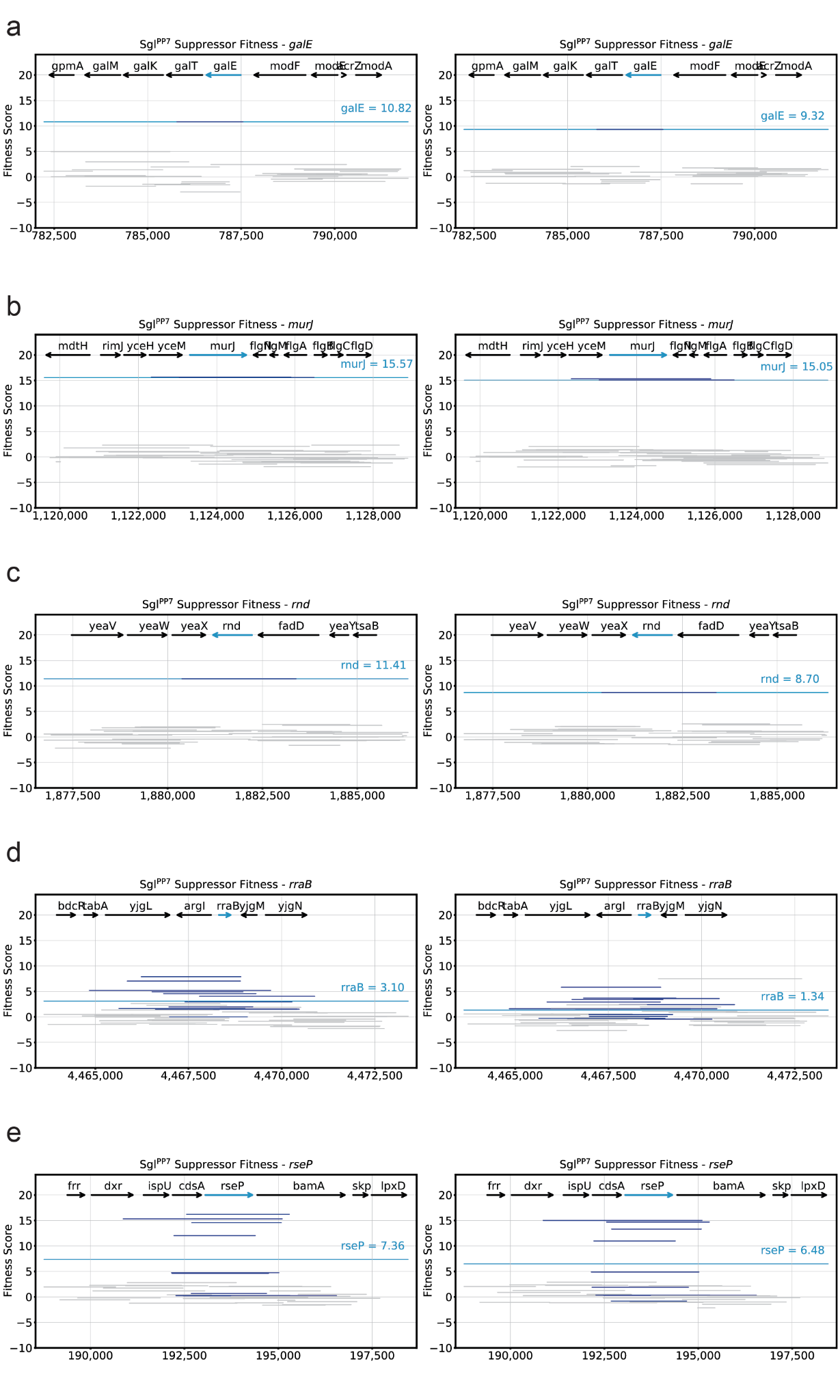

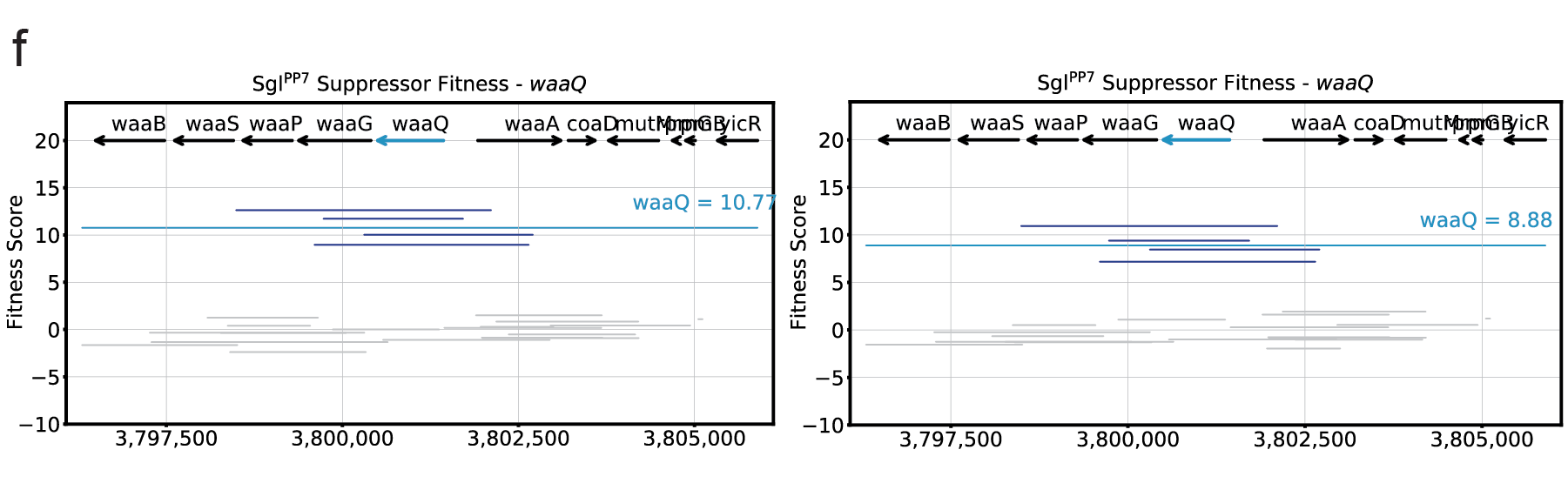

**Supplementary Fig. 5: Detailed Dub-seq suppressor screening results against Sgl^PP7^.**

(a-f) Dub-seq plots for suppressor screening against Sgl^PP7^ zoomed in on a gene of interest with experimental replicates shown side-by-side. Genes of interest are significant hits shown in Fig. 2 or candidate multi-gene multi-copy suppressors. Black arrows represent tracked ORFs during analysis. Teal arrows represent the gene of interest. Dark blue lines represent Dub-seq fragments covering the gene of interest with scores shown as *fscores*. Gray lines represent Dub-seq fragments that do not entirely cover the gene of interest with scores shown as *fscores*. Teal lines represent the *gscore* of the gene of interest.

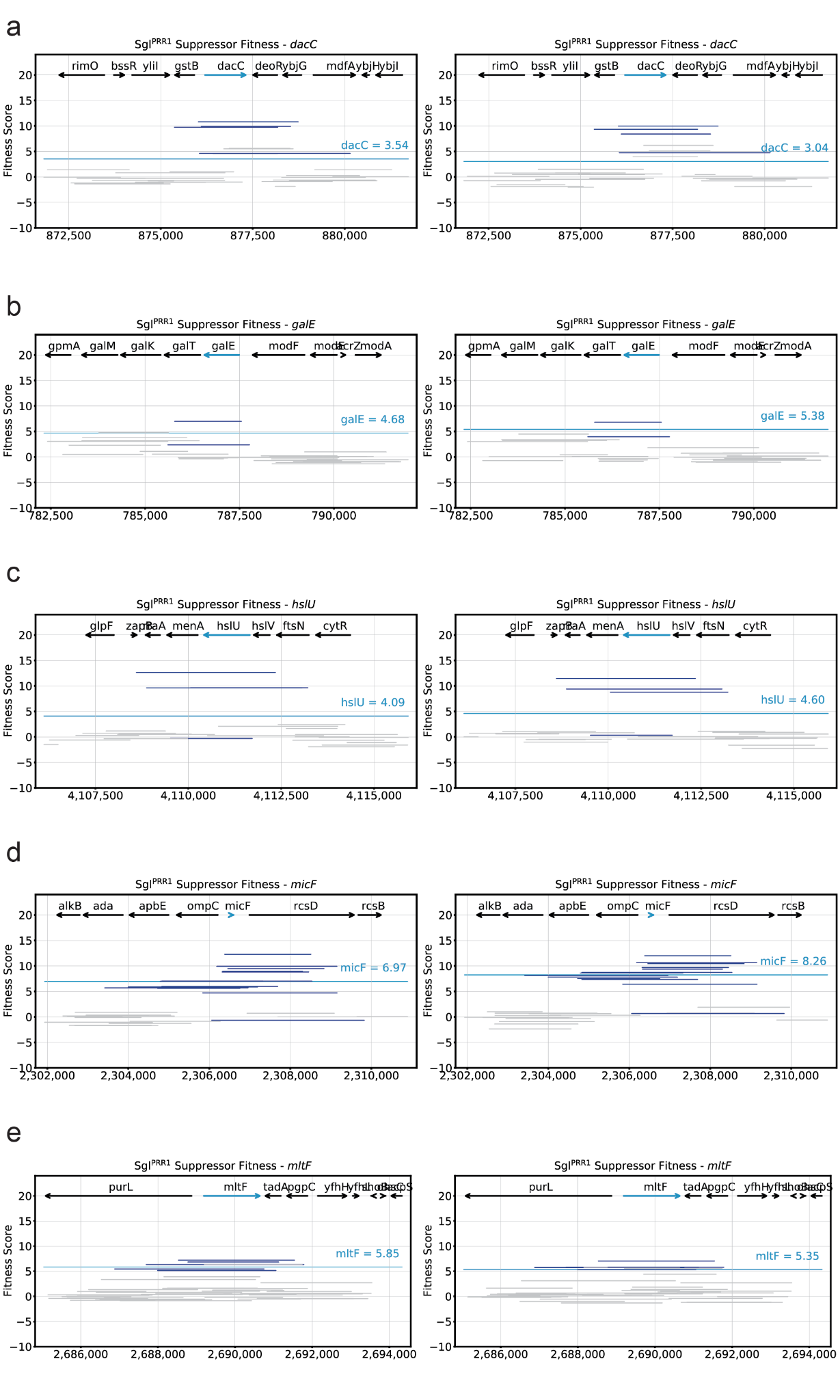

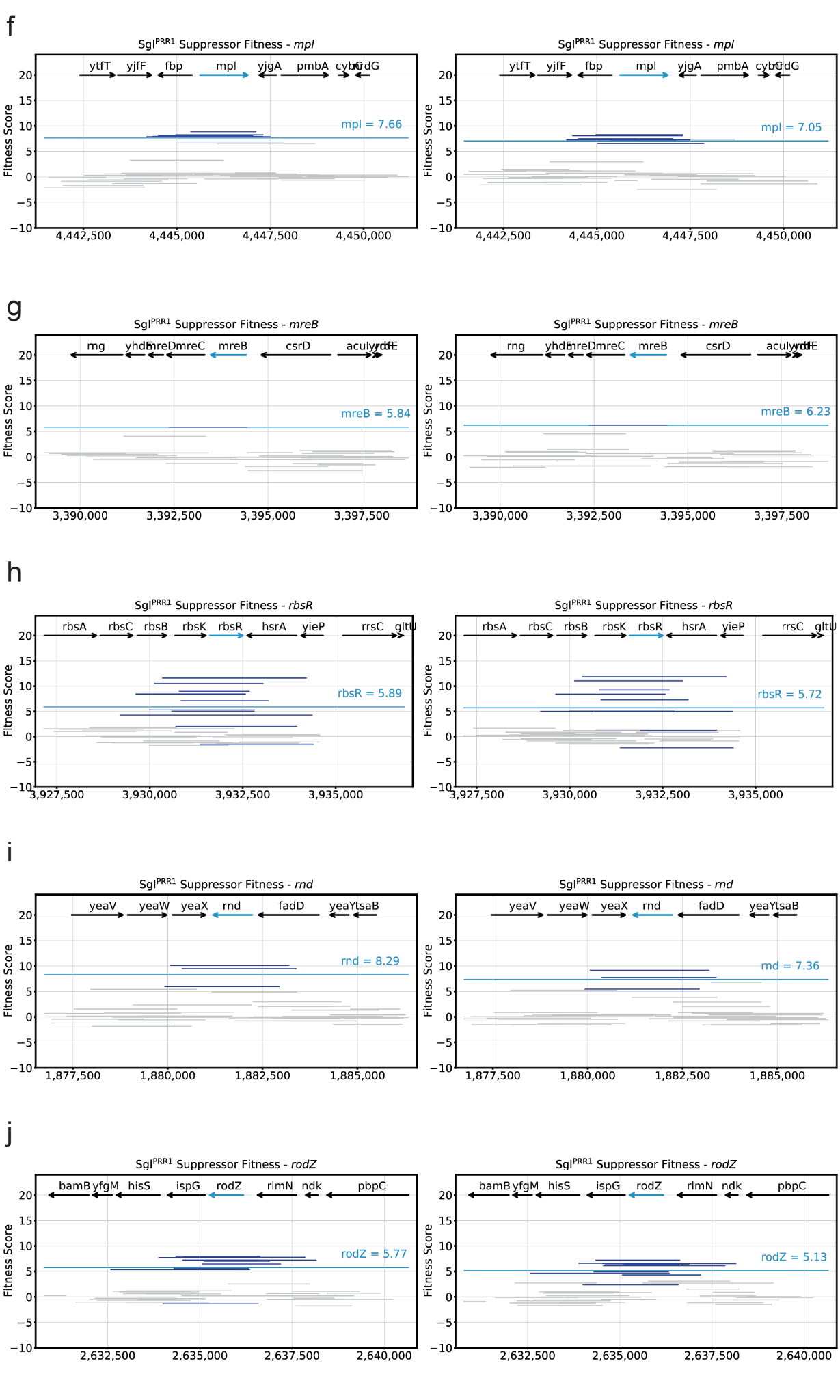

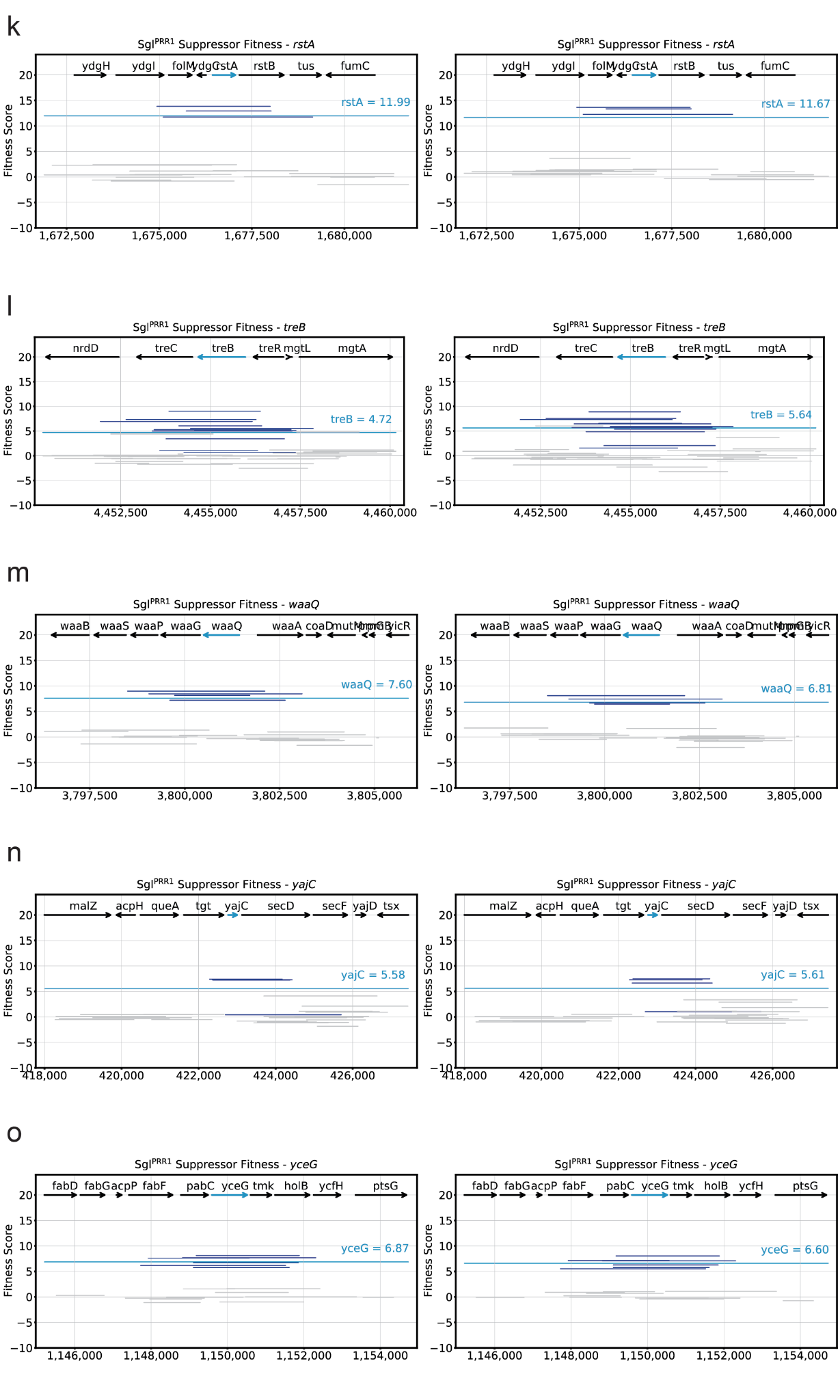

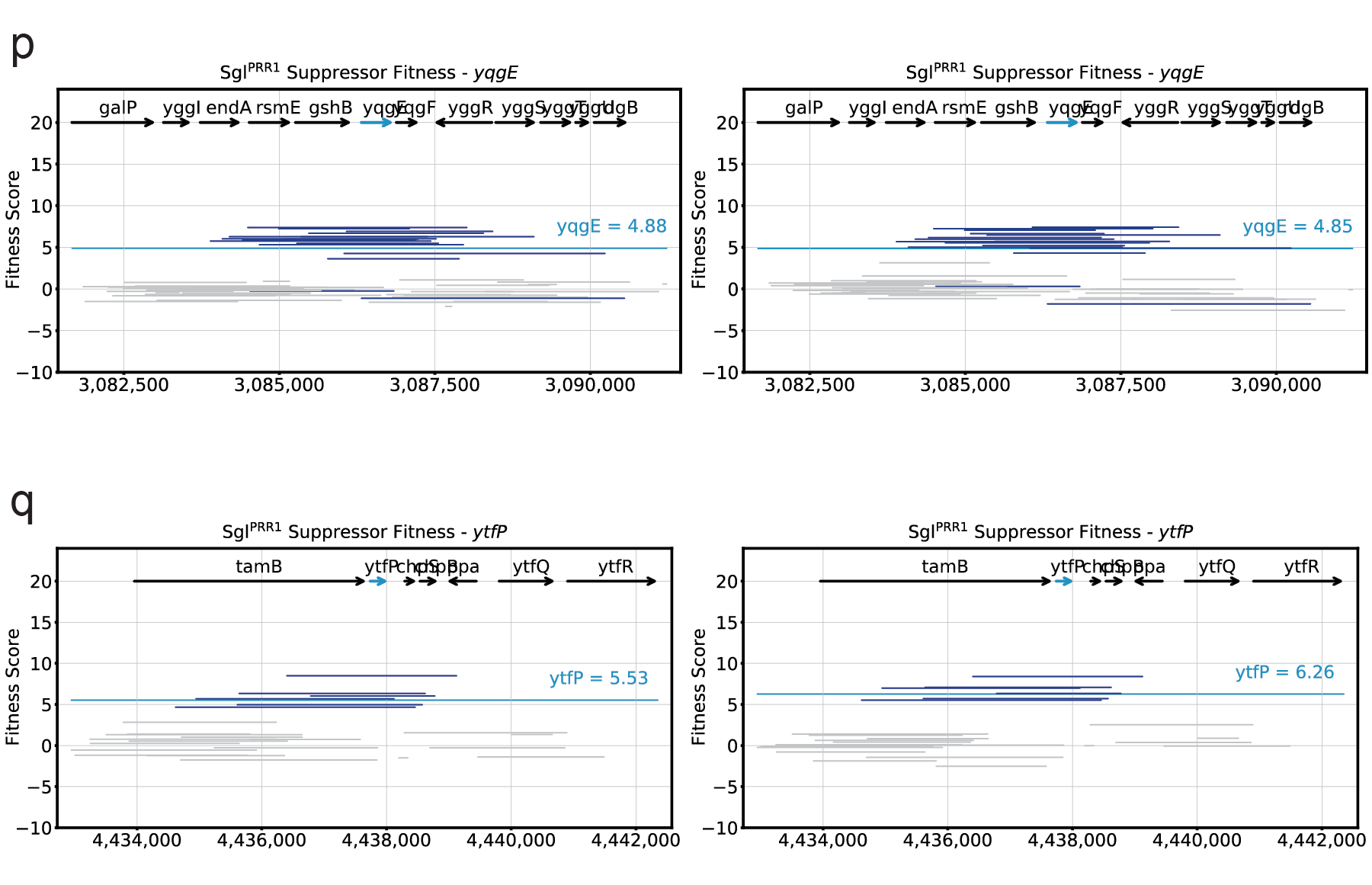

**Supplementary Fig. 6: Detailed Dub-seq suppressor screening results against Sgl^PRR1^.**

(a-q) Dub-seq plots for suppressor screening against Sgl^PRR1^ zoomed in on a gene of interest with experimental replicates shown side-by-side. Genes of interest are significant hits shown in Fig. 2 or candidate multi-gene multi-copy suppressors. Black arrows represent tracked ORFs during analysis. Teal arrows represent the gene of interest. Dark blue lines represent Dub-seq fragments covering the gene of interest with scores shown as *fscores*. Gray lines represent Dub-seq fragments that do not entirely cover the gene of interest with scores shown as *fscores*. Teal lines represent the *gscore* of the gene of interest.

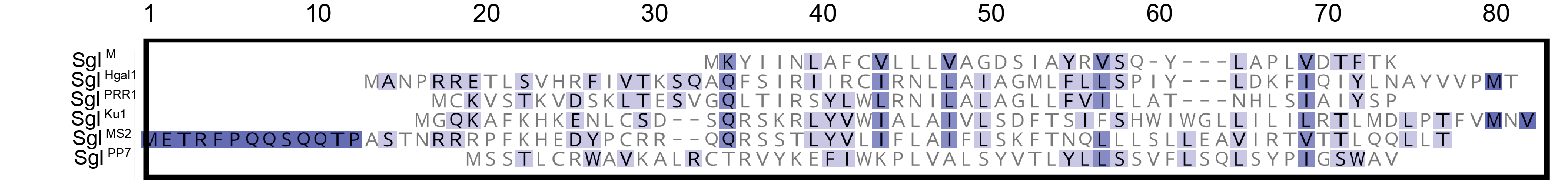

**Supplementary Fig. 7: Sequence alignment of *Fiersviridae* lysis proteins**. *Fiersviridae* lysis proteins bear little resemblance to each other. Sequence alignments were performed with MUSCLE using a BLOSUM62 matrix (PMID: 15034147) and shaded with increasing sequence similarity.

1. Recent identification and expansion of Sgls necessitates a more systematic nomenclature for these proteins, but some have less informative common nomenclature. Here, we use a format consistent with Chamakura 2020. For instance, Sgl from phage Ku1 is Sgl^Ku1^ and Sgl L from MS2 is Sgl^MS2^. In this table, the common name is provided as well.
